# Chronic industrial perturbation and seasonal change induces shift in the bacterial community from gammaproteobacteria to betaproteobacteria at Amlakhadi canal

**DOI:** 10.1101/2023.02.20.529212

**Authors:** Jenny Johnson, Kunal R. Jain, Anand Patel, Nidhi Parmar, Chaitanya Joshi, Datta Madamwar

## Abstract

Escalating proportions of industrially contaminated sites are one of the major catastrophes faced at the present time due to the industrial revolution. In the outlook of the obstacles associated with culturing the microbes, the direct metagenomic analysis of various complex niches is rapidly gaining attention. In this study, metagenomic approach using next generation sequencing technologies was applied to exemplify the taxonomic abundance and metabolic potential of the microbial community residing in Amlakhadi canal, Ankleshwar at two different seasons. All the metagenomes revealed a predominance of Proteobacteria phylum. However, difference was observed within class level where Gammaproteobacteria was relatively high in polluted metagenome in Summer while in Monsoon the abundance shifted to Betaproteobacteria. Similarly, significant statistical differences were obtained while comparing the genera amongst contaminated sites where *Serratia, Achromobacter, Stenotrophomonas* and *Pseudomonas* were abundant at one season and the dominance changed to *Thiobacillus, Thauera, Acidovorax, Nitrosomonas, Sulfuricurvum, Novosphingobium, Hyphomonas* and *Geobacter* at the next season. Further upon functional characterization, the microbiomes demonstrated diverse survival mechanisms adapted by the inherent microbial community such as degradation of aromatic compounds, heavy metal resistance, oxidative stress responses and multidrug resistance efflux pumps. The results have important implications in understanding and predicting the impacts of human-induced activities on microbial communities inhabiting natural niche and their responses in coping with the fluctuating pollution load.

## 1.0 Introduction

Anthropogenic contamination is often directed by the avid community desires and has become a serious matter of concern recurrently encountered near long-term industrialized areas. The western state of India, Gujarat, houses majority of the industrial estates within the “Golden Corridor” extending from Mehsana to Vapi (Shah et al. 2013; Sharaff et al. 2017). This industrial belt falls into the basins of various river bodies across the state making them contaminated and unhealthy for the surrounding residential areas. Consequently, the sediments act as repositories of various pollutants and xenobiotic compounds that are retained within these water bodies with time. Since the last two decades, there is an escalating awareness in the field of bioremediation due to the limitation in landfills and the increasing remediation costs. However, the successful development of any bioremediation strategy depends on understanding the inherent microbial community dynamics, structure and function (Desai et al. 2010) through genomic approaches which will further aid in comprehensive knowledge in elucidating the evolutionary history, functional and ecological biodiversity (Shokralla et al. 2012).

Metagenomic sequencing of the environmental DNA has conceded across the limitations of rRNA gene-based characterization and provided a robust platform for accessing the taxonomic affiliation and deciphering the genetic and molecular mechanism pertaining to the complex degradation processes in contaminated sites. Moreover, the latest development in next generation sequencing (NGS) technologies has facilitated many researchers to elucidate nearly complete genomes from complex niches (Hess et al. 2011; Iverson et al. 2012). Henceforth, analyzing the microbial communities through such high throughput platforms not only enable us to signify their function in a particular ecosystem but also help in statistical quantitative comparisons in community structure that happen due to fluctuations in the anthropogenic conditions (Leininger et al. 2006; Fierer et al. 2007).

It has been well studied that the innate microorganisms adapt to a diverse range of catabolic strategies to survive amidst the variety of toxic compounds prevalent in their habitat (Ufarte et al. 2015). In the course of such microbial degradation processes, the molecular structure of the xenobiotic compounds is altered chiefly due to the enzymes translated in response to the fluctuating pollution, whose specificity can accommodate the analogs of several molecules. Furthermore, with the increasing knowledge on these catabolic enzymes, their substrate efficiency to a particular compound can be modified and thus aid in discovering novel biocatalysts (Pieper and Reineke 2000, Alcalde et al. 2006). Additionally, microbes develop various evolutionary strategies to cope with the pollution load and this can direct to evolve different pathway adaptations for relating their metabolic resilience to the environment (Gianoulis et al. 2009).

In the present study, we used whole-genome shotgun (WGS) sequencing to investigate the taxonomic profile and functional diversity of the microbial community inhabiting an industrially contaminated site across space and time, which to our knowledge has not been previously explored. According to our hypothesis, the diversity in this particular niche is altered by the anthropogenic turbulence in the canal and the dissimilarities is reflected by the microbial communities and their metabolic potential to co-exist in such contaminated environment.

## 2.0 Materials and Methods

### 2.1 Site description, soil sampling and its characterization

Amlakhadi canal receives the effluents from Ankleshwar, Panoli and Jhagadia industrial estates and finally merges with Narmada. The subsurface sediment samples (5 – 8 inches below the surface) were collected in triplicate for two time points over the year (summer and monsoon of 2013) from three distinct sites (21°37’17.10”N, 72°58’56.56”E; 21°37’20.07”N, 72°58’20.15”E and 21°37’53.55”N, 72°57’23.83”E) of Amlakhadi canal. Similarly, the pristine soil samples (within 1000 m radius) without any visible pollution were collected. All the samples were immediately processed for the extraction of metagenomic DNA. The sediment samples were dried and subjected to physico-chemical characterization at Gujarat Institute of Desert Ecology, Kutch. The culturable bacterial community were analyzed as described in Supplementary information.

### 2.2 Metagenome extraction and sequencing

Metagenomic DNA from polluted and pristine sites and from each replicate was extracted independently by the method described in Desai and Madamwar (2007). The extracted DNA was purified by silica-based gel extraction kit (Merck Biosciences, Bangalore, India) as per the manufacturer’s instruction. The study was aimed to recognize the change in microbial community between polluted sites and pristine environment (i.e. inter-site difference) and functional ability of the community for sustaining the biogeography of the polluted ecosystem. Since, we did not intend to study the diversity of microbial community of individual location the metagenome extracted from each polluted site (for each time point separately) was mixed in equal concentration to make a composite DNA, representing the polluted environment. Likewise, the extracted DNA from the pristine soils were also mixed in equal concentration to obtain the composite metagenomic DNA. These composite DNA were sequenced on Ion Torrent Platform using 316 chip with 200 bp chemistry.

### 2.3 Taxonomic and functional annotation of metagenomes

The sequenced datasets were deposited in MG-RAST server version 3.5 (Meyer et al. 2008) for taxonomic and functional annotation of the sequences. Artificial replicate sequences were removed according to Gomez-Alvarez et al. (2009) and low quality sequences (quality mean value 20%) were removed using PRINSEQ (Schmieder and Edwards 2011). The taxonomic profiles were analysed using M5NR database, while the reads for functional genes were annotated using SEED subsystem database (Overbeek et al. 2005). The best hit classification at a maximum e-value of 1e^−5^, a minimum identity of 60% and a minimum alignment length of 15 were used in both the analysis. For identification of genes, the functions and their corresponding taxa, the open reading frames (ORF) were predicted from the sequenced reads using FragGeneScan (Rho et al. 2010). These ORF’s were annotated against the KEGG database to characterize the metabolic pathways and GhostKOALA (Kanehisa et al. 2016) to identify the taxonomy associated with the particular functions in the specific pathway.

### 2.4 Statistical analysis

The statistical comparison between the taxonomic and functional profiles were carried out by using the Statistical Analyses of Metagenomic Profiles (STAMP) software v 2.1.3 (Parks et al. 2014). Significant statistical differences between samples were processed by two-sided Fisher’s exact test with Newcombe-Wilson confidence interval method, and Benjamini-Hochberg FDR and results with p values (p<0.05) were considered. The diversity indices were computed using PAST Version 3.06 software using default parameters.

### 2.5 Sequence Accession Numbers

All metagenome sequences have been deposited to the MG-RAST database under the accession numbers 4541801.3 (Polluted 1), 4542275.3 (Control 1), 4618428.3 (Polluted 2) and 4607054.3 (Control 2). The 16S rRNA gene sequences from the culturable bacterial community were submitted to NCBI with Accession No. KX817810 to KX817971.

## 3.0 Results

The polluted and control soil samples collected during summer (Season 1) and monsoon (Season 2) of the year 2013 were subjected to high throughput sequencing using Ion Torrent PGM platform. The metagenomes from Season 1 were designated as P1 for polluted and C1 for control while for Season 2, P2 for polluted and C2 for control samples. The physico-chemical characteristics of the polluted and control soil samples collected from Amlakhadi canal are tabulated in Supplementary Table S1. The pH of the soil samples ranged from 7.8-8.4 indicating slightly alkaline conditions. The electrical conductivity of the polluted soil varied between 16.6 MS - 48.1 MS while in the control soil it was 10.1 MS. Heavy metals such as copper, manganese and zinc were found in elevated amounts contributing in many industrial processing activities.

### 3.1 Microbial Community Structure deciphered by shotgun sequencing

The classification of microbial lineage was generated by annotating the sequenced reads against the M5NR database using the parameters described above. According to the classifier, the Bacterial domain was most dominant in all the four metagenomes i.e. P1 (98.6%), P2 (95.3%), C1 (96.6%) and C2 (98.9%). In phylum classification, it was characterized that the major dominant phylum in all the metagenomes was Proteobacteria (84.6% - 98.6%) as shown in Figure 1, which was followed by a minor abundance of Actinobacteria in P1 (1.7%), P2 (0.94%) and C1 (11.2%) whereas Firmicutes was the next abundant phyla in C2 (0.52%). At rank class, in polluted samples, the abundant bacterial class in P1 metagenome was Gammaproteobacteria (48.5%) whereas at the next season, the abundance shifted to Betaproteobacteria (46.6%) in P2 metagenome. In P1 metagenome, the next dominant class was Betaproteobacteria (26.2%) followed by Alphaproteobacteria (16.8%) and Deltaproteobacteria (3.7%). During the monsoon, the second abundant class in P2 metagenome was Gammaproteobacteria (19.6%) followed by Alphaproteobacteria (18.1%) and Epsilonproteobacteria (6.6%). Whilst in control samples, both the seasonal metagenome showed the abundance of Gammaproteobacteria in C1 (62.8%) and C2 (98.7%) respectively which was followed by Betaproteobacteria (18.5%) and Actinobacteria (11.3%) in C1 and Bacilli (0.38%) in C2 metagenome.

**Figure 1.**
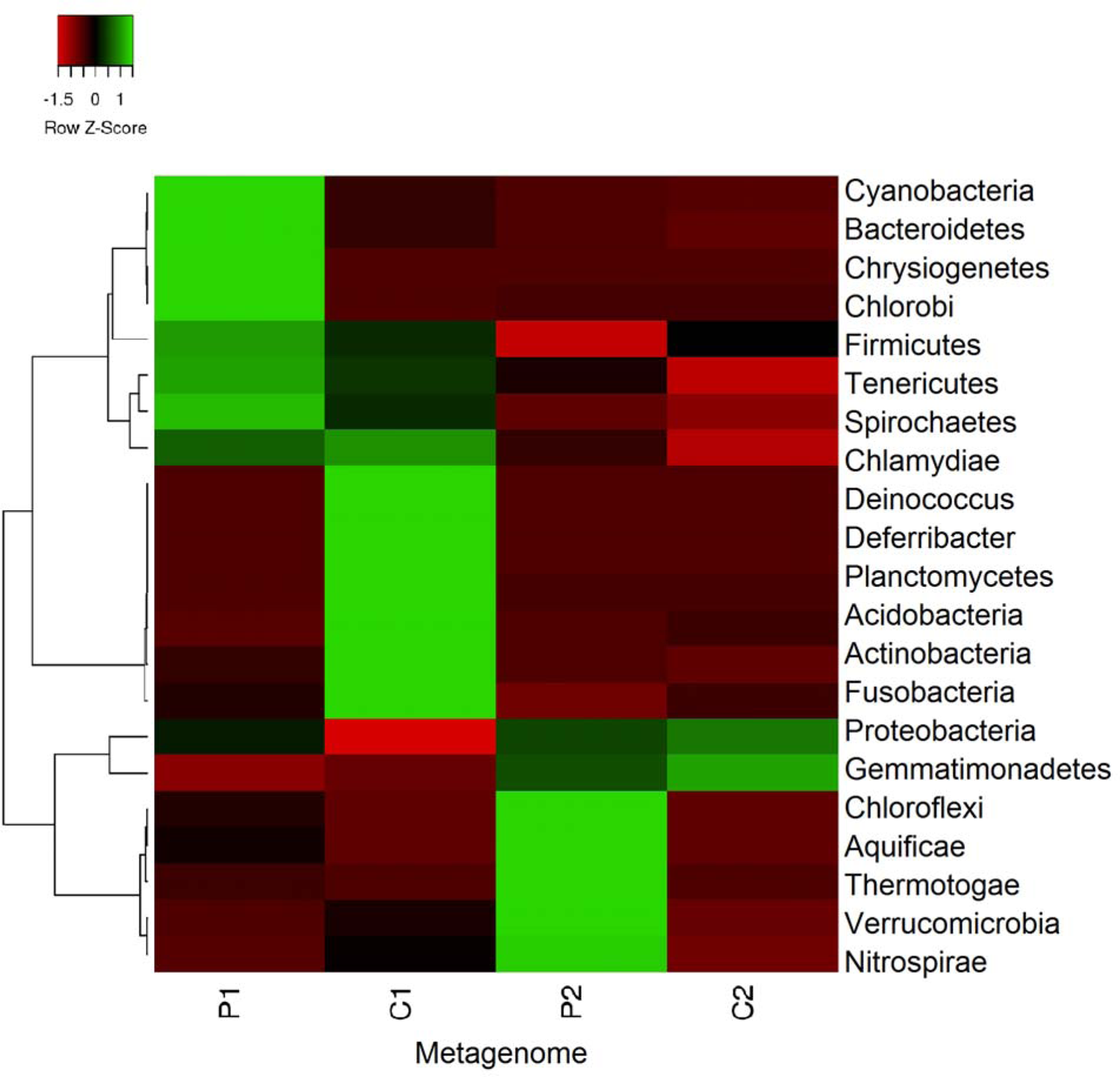
Heatmap representing the phylum level composition in P1, C1, P2 and C2 metagenome annotated against M5NR database in MGRAST server with 60% identity and e-value of 10^-5^.

Comparative statistical analysis of the metagenomes revealed significant differences at genus level between both the seasons. During Season 1 (summer), the results demonstrated that *Serratia* (37.2%), *Achromobacter* (11.5%) and *Stenotrophomonas* (9.7%) were the genus that were abundant in C1 metagenome when statistically analyzed (p<0.05) (Figure S1). Surprisingly, the P1 metagenome also showed the abundance of *Serratia* (22.9%), *Achromobacter* (7.1%) and *Stenotrophomonas* (4.7%) which was followed by *Pseudomonas* (3.1%), *Yersinia* (2.8%), *Bordetella* (2.7%), *Xanthomonas* (1.8%), *Acidovorax* (1.7%), *Burkholderia* (1.6%) and *Thiobacillus* (1.3%) that were statistically significant (*p* < 0.05). Whereas in the next season (monsoon), the significant statistic difference (*p* < 0.05) was observed between 30 different genera in P2 and C2 metagenome (Figure S2). *Serratia* (73%), *Stenotrophomonas* (7.3%), *Yersinia* (4.1%) and *Escherichia* (2.1%) were in abundance in control (C2) metagenome. Whereas *Thiobacillus* (16.4%), *Thauera* (8.6%), *Acidovorax* (7.93%), *Nitrosomonas* (5.4%), *Sulfuricurvum* (4.8%), *Novosphingobium* (4.6%), *Hyphomonas* (3.6%), *Geobacter* (3.3%), *Sphingobium* (3%), *Alicycliphilus* (2.9%), *Legionella* (2.4%), *Pseudomonas* (2.4%), *Oligotropha* (2.3%) and *Desulfovibrio* (2.2%) were some of the dominant genera scrutinized in P2 metagenome. Of the 480 genera identified amongst all the metagenomes, we compared to recreate a Venn diagram (Figure S3) to identify the potential rare phylotypes within each metagenome and it is tabulated in Supplementary Table S2. By analyzing the taxonomic profile through culturable approach, both the contaminated and control soil, were dominated by Firmicutes. *Bacillus* genus were found in abundance (77-80%) in both the samples (Figure S4).

#### 3.1.1. Diversity indices

To understand the extent of diversity within the samples, diversity indices for all the samples was computed using PAST3 (Table S3) and Principal component analysis (PCA) plot was generated (Figure S5). Higher diversity as indicated from Chao, Fisher_Alpha and Shannon indices was observed in P1, P2 and C1 while the lowest diversity was observed in C2. Simpson index values exceeded 0.9 for both the polluted metagenome whereas the control samples showed the value of 0.8 (C1) and 0.4 (C2) respectively. Dominance_D was high in C2 (0.5) when compared to rest of the samples.

### 3.2 Functional metabolic profiling of the inherent community

Functional distributions were characterized by SEED subsystems wherein out of the 28 subsystems in level 1, both the polluted metagenomes (P1, P2) were abundant in clustering based subsystems (∼14%) whereas in control metagenomes (C1, C2) carbohydrates (∼16%) was the dominant which was followed by amino acids and derivatives (9-10%) and protein metabolism (6-9%) (Figure S6). Considering the perturbed environment, the subsystems like metabolism of aromatic compounds, stress responses and virulence, disease and defense were further deciphered at Level 2. All the metabolic profiles were statistically analyzed in STAMP and the results stated are significantly different (*p* < 0.05).

#### 3.2.1 Metabolism of aromatic compounds

As seen in Figure 2, the genes corresponding to n-Phenylalkanoic acid degradation were the prominent in P1 (24.2%), P2 (42.3%) and C1 (19.2%). Phenylacetyl-CoA catabolic pathway (8.8%) and Benzoate transport and degradation cluster (2.6%) were relatively abundant in P1 metagenome when compared with C1. In season 1, salicylate and gentisate catabolism (7.3%) and N-heterocyclic aromatic compound degradation (3.3%) were significantly higher (p<0.05) in control metagenome (C1) than the polluted metagenome (P1) (Figure 2a) whereas in season 2, genes involved in 4-Hydroxyphenylacetic acid catabolic pathway (13.7%), Phenylacetyl-CoA catabolic pathway (7.9%), Aromatic Amin Catabolism (5.8%) and Hydroxyaromatic decarboxylase family (1.6%) were in elevated levels in control metagenome (C2) than polluted metagenome (P2) (Figure 2b).

**Figure 2.**
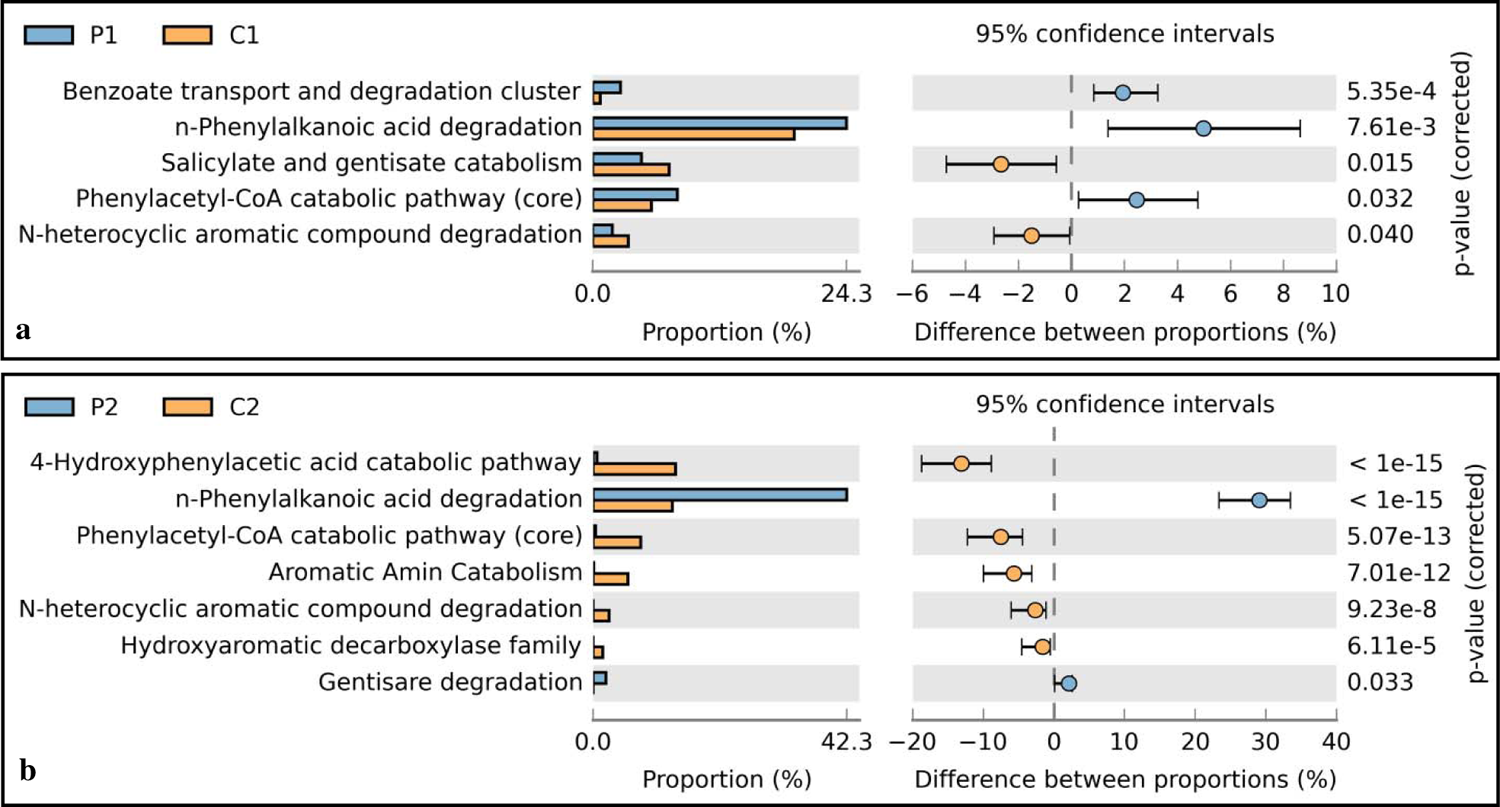
Statistically significant differences in the functional subsystem Metabolism of Aromatic compounds of SEED classification using STAMP statistical software in the comparison of (a) P1 and C1 and (b) P2 and C2. The graphic shows only the subsystems with statistical differences between the proportions of sequences in each metagenome with a confidence interval of 95%.

#### 3.2.2 Stress responses

Anthropogenic conditions often induce stress in the inherent microbial community which was quite evident in the Stress responses functional category annotated by SEED subsystems (Figure 3). Increased levels of genes involved in regulation of oxidative stress response were observed in the metagenomes specifically in C1 (16%), P1 (12.9%) (Figure 3a) and C2 (6.47%) while quite low level of 1.3% in P2. However, P2 metagenome was enriched with genes related to Protection from Reactive oxygen species (8.5%), Redox dependent regulation of nucleus processes (8.3%), Rubrerythrin (6.4%), Bacterial hemoglobins (5.4%) and Acid resistance mechanisms (2.9%) (Figure 3b).

**Figure 3.**
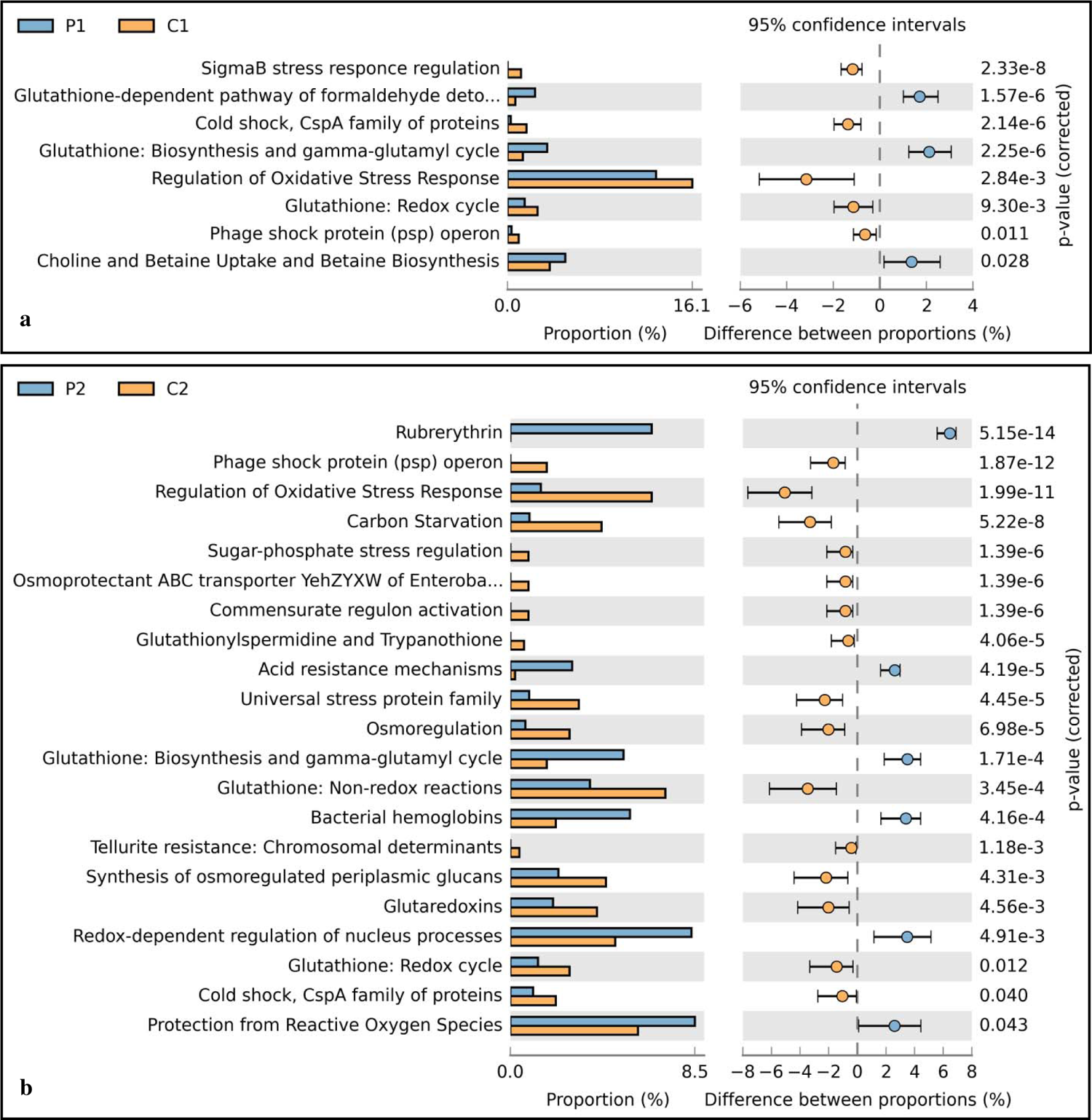
Statistically significant differences in the functional subsystem Stress Responses of SEED classification using STAMP statistical software in the comparison of (a) P1 and C1 and (b) P2 and C2. The graphic shows only the subsystems with statistical differences between the proportions of sequences in each metagenome with a confidence interval of 95%.

#### 3.2.3 Virulence, Disease and Defense

Majority of the sequenced reads from all the metagenomes were annotated in the Virulence, Disease and Defense category of SEED (Figure 4). Cobalt-zinc-cadmium resistance (10-25%), Copper Homeostasis (7-8%), Arsenic resistance (3-7%), Mercury resistance operon (0.5-2.5%), mercuric reductase (0.4-2%) and beta lactamases (0.5-3%) were prevalent abundantly in polluted metagenome P1 and P2. The control metagenome, C1 and C2, showed the abundance of genes pertaining to multidrug resistance efflux pumps (15-18%), resistance to fluroquinolones (13%), MLST (12%), bacterial cyanide production and tolerance mechanisms (3-9%).

**Figure 4.**
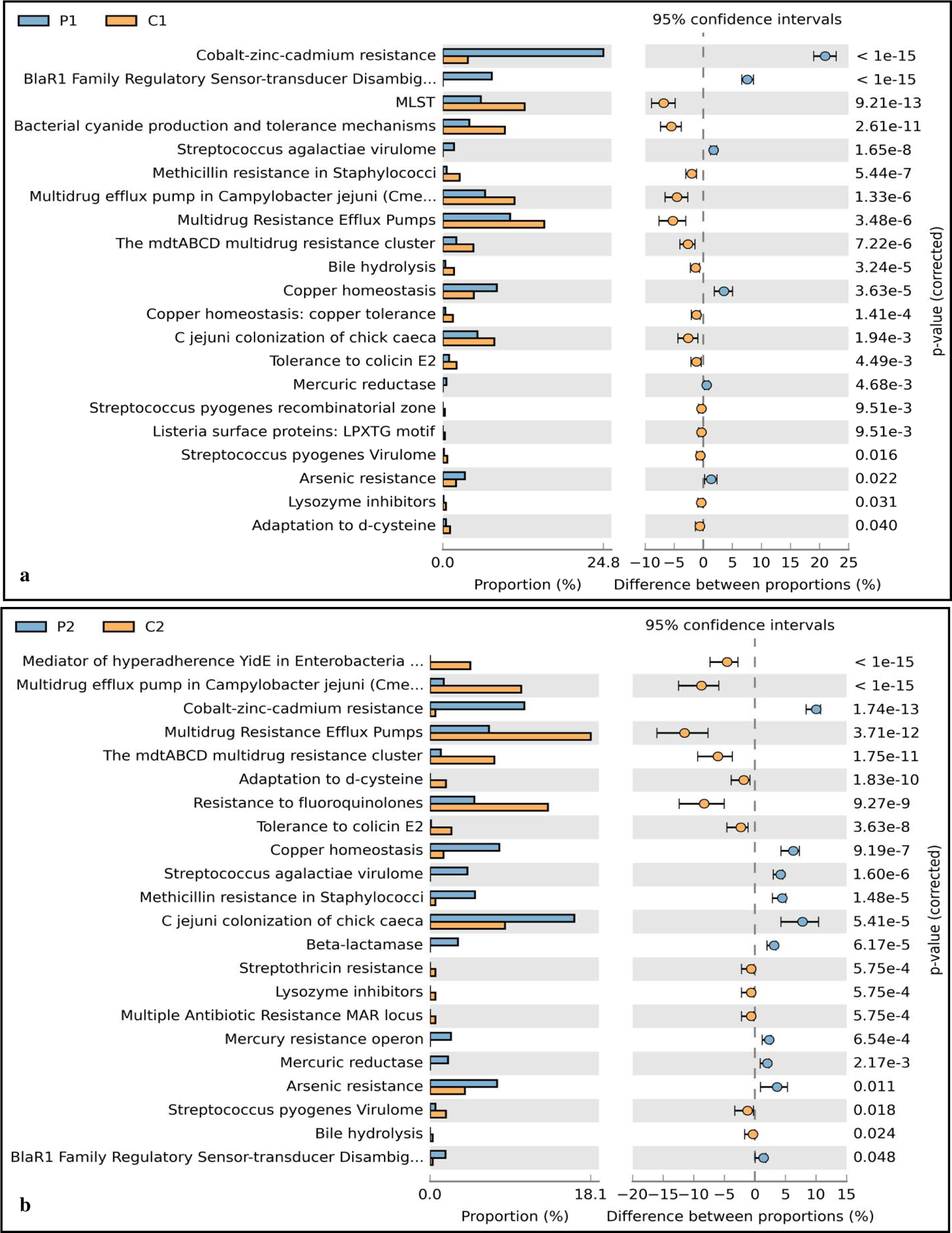
Statistically significant differences in the functional subsystem Virulence, Disease and Defense of SEED classification using STAMP statistical software in the comparison of (a) P1 and C1 and (b) P2 and C2. The graphic shows only the subsystems with statistical differences between the proportions of sequences in each metagenome with a confidence interval of 95%.

#### 3.2.4 Corelating the metagenomes by functional categories of SEED

The Level 3 classification of SEED subsystems was used to co-relate the functional genes between the samples based on their R^2^ values generated in the scatter plots in STAMP. As evident in Figure 5, there was a higher correlation (R^2^=0.836) between P1 and C1 (Figure 5a) and least correlation (R^2^=0.332) between P2 and C2 (Figure 5b). Comparing the similar profiles, more correlation (R^2^=0.735) was obtained between C1 and C2 (Figure 5c) rather than P1 and P2 (Figure 5d) having a weaker correlation of R^2^=0.419.

**Figure 5.**
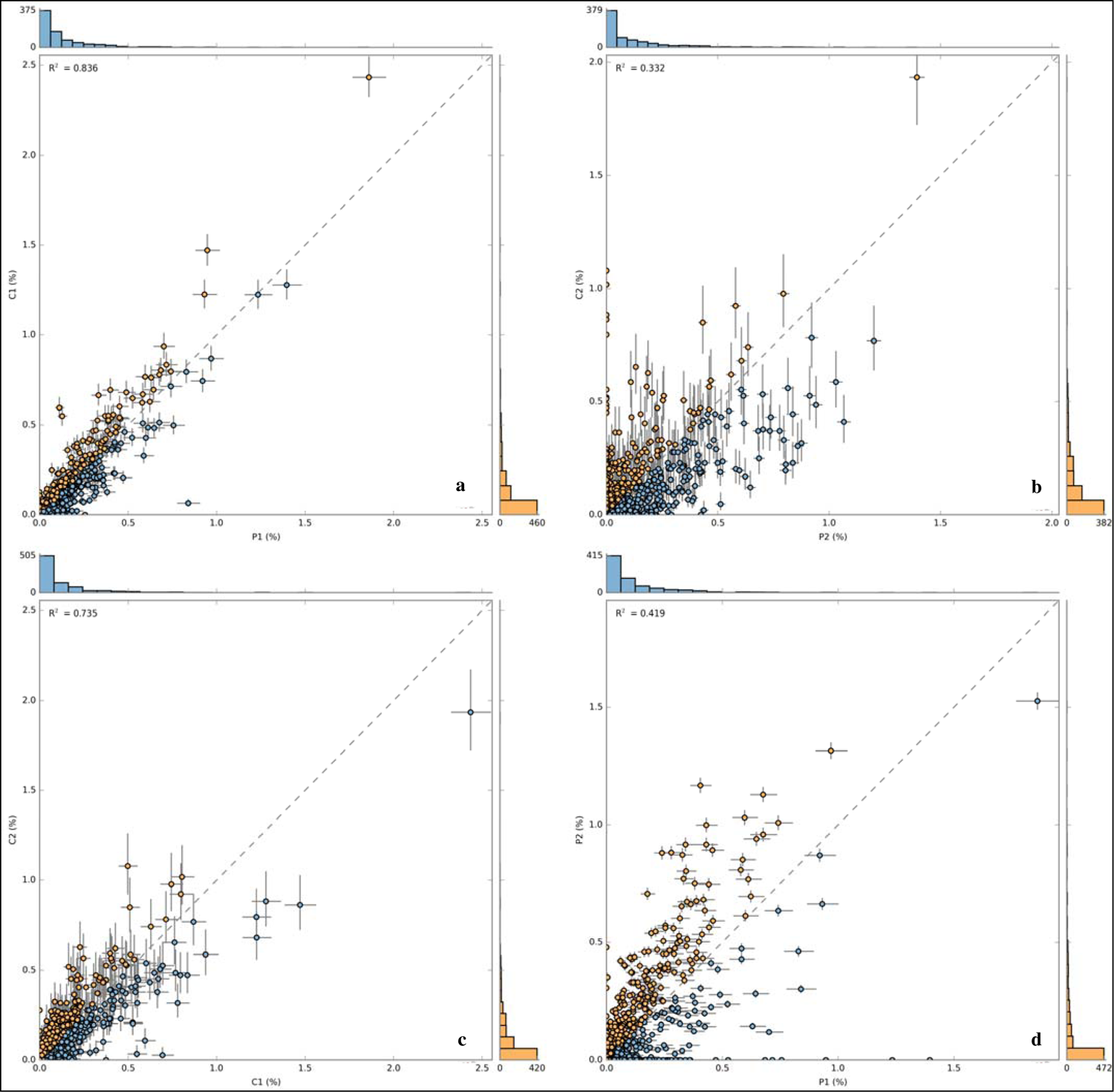
Scatter plots of functional gene annotations using SEED subsystem level III of (a) P1 and C1 (b) P2 and C2 (c) C1 and C2 (d) P1 and P2. Each dot represents a unique functional classification gene.

#### 3.2.5 KEGG pathway analysis

A detailed analysis of KEGG pathways was carried out based on the KEGG orthology (KO) of all the four metagenomes. Since we were essentially paying attention to the metabolism pathways of xenobiotics biodegradation, it was found that enzymes involved in this category were relatively abundant (835 hits) in the bacterial metagenomes. Among them, the genes encoding enzymes involved in the pathway of benzoate degradation were more abundant (21-24%) than in any other pathways (Figure S7), with a complete elucidation of catechol breakdown to acetyl-CoA in the benzoate degradation via hydroxylation pathway (Figure S8). Similarly, majority of the enzymes were mapped for aminobenzoate (Figure S9) and toluene and xylene degradation (Figure S10) in both the polluted and control metagenomes. Almost 40 different enzymes were mapped from the metagenome reads catabolizing in these three degradation pathways and their abundance in each metagenome are tabulated in Table 1. Carboxymethylenebutenolidase, 4-hydroxy 2-oxovalerate aldolase, 4-carboxymuconolactone decarboxylase, 4-oxalmesaconate hydratase, 4-oxalocrotonate tautomerase, 2-hydroxy-4-carboxymuconate semialdehyde hemiacetal dehydrogenase, 3-oxoadipate CoA-transferase, alpha subunit, nitrile hydratase subunit alpha, phenol 2-monooxygenase, p-hydroxybenzoate 3-monooxygenase, catechol 2,3-dioxygenase were some of the abundant enzymes found in the metagenome P1, C1 and P2 whilst C2 metagenome showed lower hits corresponding to these enzymes. The predicted ORF’s from the metagenome were analyzed in GhostKOALA to assign their taxonomic affiliation to the functions associated with KEGG pathways. The enzymes that were mapped in the degradation of xenobiotics were further scrutinized to identify their phylogenetic affiliation. The genera to the respective enzyme was identified based on their GhostX score in GhostKOALA web server and are tabulated in Supplementary Table S4.

**Table 1.** List of enzymes involved in the benzoate, aminobenzoate, toluene and xylene degradation pathways using KEGG database in KEGG orthology and their corresponding number of hits identified in each metagenome.

| Enzymes | Metagenome |  |  |  |
| --- | --- | --- | --- | --- |
|  | P1 | C1 | P2 | C2 |
| carboxymethylenebutenolidase [EC:3.1.1.45] | 24 | 69 | 129 | 19 |
| catechol 2,3-dioxygenase [EC:1.13.11.2] | 5 | 4 | 19 | 7 |
| 4-oxalomesaconate hydratase [EC:4.2.1.83] | 7 | 24 | 50 | 7 |
| 3-oxoadipate CoA-transferase, alpha subunit [EC:2.8.3.6] | 4 | 3 | 27 | 6 |
| muconate cycloisomerase [EC:5.5.1.1] | 5 | 3 | 8 | 5 |
| nitrile hydratase subunit alpha [EC:4.2.1.84] | 5 | 16 | 25 | 4 |
| protocatechuate 3,4-dioxygenase, alpha subunit [EC:1.13.11.3] | 4 | 5 | 10 | 4 |
| catechol 1,2-dioxygenase [EC:1.13.11.1] | 2 | 8 | 15 | 3 |
| 4-hydroxy 2-oxovalerate aldolase [EC:4.1.3.39] | 18 | 13 | 71 | 3 |
| benzoylformate decarboxylase [EC:4.1.1.7] | 0 | 11 | 12 | 3 |
| acetyl-CoA acyltransferase 2 [EC:2.3.1.16] | 0 | 2 | 3 | 3 |
| dihydroxycyclohexadiene carboxylate dehydrogenase [EC:1.3.1.25 1.3.1.-] | 1 | 0 | 5 | 3 |
| benzoate/toluene 1,2-dioxygenase subunit beta [EC:1.14.12.10 1.14.12.-] | 4 | 3 | 1 | 3 |
| pentachlorophenol monooxygenase [EC:1.14.13.50] | 0 | 2 | 3 | 2 |
| aryl-alcohol dehydrogenase [EC:1.1.1.90] | 2 | 4 | 12 | 2 |
| p-hydroxybenzoate 3-monooxygenase [EC:1.14.13.2] | 9 | 24 | 20 | 2 |
| 4-carboxymuconolactone decarboxylase [EC:4.1.1.44] | 13 | 25 | 64 | 2 |
| mandelate racemase [EC:5.1.2.2] | 0 | 1 | 1 | 1 |
| phenol 2-monooxygenase [EC:1.14.13.7] | 5 | 6 | 24 | 1 |
| 4-oxalocrotonate tautomerase [EC:5.3.2.6] | 1 | 4 | 40 | 1 |
| 3-oxoadipate enol-lactonase [EC:3.1.1.24] | 5 | 5 | 9 | 1 |
| 5-carboxymethyl-2-hydroxymuconate isomerase [EC:5.3.3.10] | 0 | 4 | 23 | 1 |
| 4-hydroxy-4-methyl-2-oxoglutarate aldolase [EC:4.1.3.17] | 0 | 1 | 12 | 1 |
| 2,4-dichlorophenol 6-monooxygenase [EC:1.14.13.20] | 0 | 2 | 8 | 0 |
| chloromuconate cycloisomerase [EC:5.5.1.7] | 0 | 0 | 1 | 0 |
| maleylacetate reductase [EC:1.3.1.32] | 0 | 1 | 10 | 0 |
| hydroxyquinol 1,2-dioxygenase [EC:1.13.11.37] | 0 | 3 | 15 | 0 |
| 2-hydroxymuconate-semialdehyde hydrolase [EC:3.7.1.9] | 1 | 0 | 0 | 0 |
| 2-keto-4-pentenoate hydratase [EC:4.2.1.80] | 0 | 4 | 8 | 0 |
| benzaldehyde dehydrogenase (NAD) [EC:1.2.1.28] | 0 | 0 | 6 | 0 |
| 2-oxo-3-hexenedioate decarboxylase [EC:4.1.1.77] | 1 | 0 | 14 | 0 |
| aminomuconate-semialdehyde/2-hydroxymuconate-6-semialdehyde dehydrogenase [EC:1.2.1.32 1.2.1.85] | 2 | 0 | 7 | 0 |
| 4-cresol dehydrogenase (hydroxylating) flavoprotein subunit [EC:1.17.99.1] | 0 | 4 | 0 | 0 |
| anthranilate 1,2-dioxygenase (deaminating, decarboxylating) large subunit [EC:1.14.12.1] | 0 | 0 | 1 | 0 |
| 3-oxoadipyl-CoA thiolase [EC:2.3.1.174] | 2 | 3 | 14 | 0 |
| muconolactone D-isomerase [EC:5.3.3.4] | 3 | 0 | 4 | 0 |
| protocatechuate 4,5-dioxygenase, alpha chain [EC:1.13.11.8] | 3 | 1 | 11 | 0 |
| 2-pyrone-4,6-dicarboxylate lactonase [EC:3.1.1.57] | 1 | 3 | 19 | 0 |
| 2-hydroxy-4-carboxymuconate semialdehyde hemiacetal dehydrogenase [EC:1.1.1.312] | 8 | 0 | 28 | 0 |
| 3-carboxy-cis,cis-muconate cycloisomerase [EC:5.5.1.2] | 4 | 13 | 5 | 0 |

### 4.0 Discussion

The present study provides the first report on the microbial diversity of the long term contaminated Amlakhadi canal, Ankleshwar using metagenomic approaches. Amlakhadi Canal, a tributary of Narmada river, stands out in being one of the five most polluted river stretches in India (CPCB 2010) which is impinged with all the toxic industrial wastes that is released into this tributary by the surrounding Ankleshwar industrial estate (Gujarat, India).

The taxonomic profile reveals that Proteobacteria phyla are widely distributed in all the four metagenomes comprising of nearly 85-98% of the total obtained reads. Proteobacteria has been the abundant phyla in many of the studies including petroleum contaminated Arctic soils (Bell et al. 2011), polluted Permafrost soils in China (Yang et al. 2012), long term diesel contaminated soil from Poland (Sutton et al. 2013), soil contaminated by heavy metals (Gołę 2014) and polluted sediments from Sea (Korlević et al. 2015). This group of bacteria is highly versatile; however, studies have proved that upon hydrocarbon contamination, there is a shift in the microbial community towards enrichment of Proteobacteria (Labbe et al. 2007; Bell et al. 2011; Zhang et al. 2012; Adetutu et al. 2013).

The genus *Serratia* has been reported in the removal of hexavalent chromium (Joutey et al. 2014), phenol and ammonium (Lu et al. 2014) and treatment of paper mill effluent (Haq et al. 2016). The fact that the three metagenomes (P1, C1, C2) showing the dominance of *Serratia* affirms that there is not much taxonomic difference between these sites suggesting that the pristine sites are contaminated due to the industrial pollution prevalent in that area for the last four decades now. *Thiobacillus* was the abundant genera in P2 sample, they are a major group of organisms belonging to potent sulphate oxidizers and encompass binding sites for dissolved metal that further serve as nucleation surfaces (Fortin et al. 1996; Torrentó et al. 2010). *Pseudomonas*, *Yersinia*, *Bordetella*, *Xanthomonas*, *Acidovorax*, *Burkholderia* were the few other genera present in the polluted sample of Summer (P1) whilst in Monsoon (P2) there was a community shift towards *Thauera*, *Acidovorax*, *Nitrosomonas*, *Sulfuricurvum*, *Novosphingobium*, *Hyphomonas*, *Geobacter*, *Sphingobium*, *Alicycliphilus*, *Legionella*, *Pseudomonas*, *Oligotropha* and *Desulfovibrio*. Various reports draw attention towards the catabolic potential of *Pseudomonas* to degrade polyaromatic hydrocarbons (Isaac et al. 2015), dye and dye effluents (Chougale et al. 2014; Garg et al. 2015). Iron oxidizers like *Acidovorax* genus contribute in PAHs degradation (Singleton et al. 2009; Patel et al. 2014). Altering the membrane fluidity upon the binding of hydrocarbon to the cell membrane to tolerate toxicity is reported in genera such as *Novosphingomonas* and *Sphingobium* (Baraniecki et al. 2002; Aylward et al. 2013). *Geobacter* is an important metal ion-reducing bacterium (Badalamenti et al. 2015) that can reduce Fe(III) (Zhou et al. 2014), Pd(II) (Pat-Espadas et al. 2013) and uranium (VI) (Prakash et al. 2010). *Desulfovibrio* genera can treat metal-containing wastewater (Karnachuk et al. 2015) with applications such as Hg methylation (Goni-Urriza et al. 2015) and uranium reduction (Payne et al. 2002). These findings indicate that there was a community shift in the polluted samples from one season to other which moved towards more tolerant genera and this shift might be due to the diverse range of effluents flowing into the canal from time to time. The community overlaps depicted in the Venn diagram (Figure S3) demonstrated a higher prevalence of some rare genera in P1(104 genus), P2 (58 genus), C1 (48 genus) whereas only 11 genera in C2. Analyzing the rare species in any ecology studies is noteworthy (Preston 1962; May 1975; Tokeshi 1993) since they have significant role in the functioning of any ecosystem and its stability that can be altered due to environmental fluctuations (MacDougall et al., 2013). Nevertheless, considering the higher abundance of common species they excessively affect the ecosystem function (Smith and Knapp 2003; Gaston 2010) but, rare genera being in relatively lower abundance cannot be ignored as such rare species can have extraordinary features or synergism with other species that sway the functioning of any niche (Lyons et al. 2005; Hooper et al. 2005; Mouillot et al. 2013). The Principal Component Analysis (PCA) represents that P2 stands out of all the metagenomes falling into distinct co-ordinate describing its unique microbial composition whereas P1 and C2 cluster together and even C1 is nearer to them which indicates that P1, C2 and C1 have somewhat similar taxonomic profiles. Diversity indices reveal higher diversity in P1, P2 and C1 metagenome as indicated from Simpson, Shannon, Chao, Fisher_alpha indices while the C2 metagenome had lower diversity comparatively and a higher dominance D level which can be due to the high abundance of only one genus (Serratia ∼ 73%). Culturable microbial diversity indicates the pre-dominance of *Bacillus* (77-80%) in both the samples whereas in the metagenomic analysis their dominance was fairly low. This difference can be attributed to the limitations of culture dependent techniques that represent only ∼ 0.1% of the entire niche (Handelsman 2004). Although the other genera were present but due to their relatively low abundance (> 2%), we cannot forecast the actual scenario of any ecosystem. Henceforth, relying only on culture dependent approaches often mislead us to the actual representation of the community structure which can be accomplished by comprehensive metagenomic studies.

Long term perturbation of any environmental niche not only affects the taxonomic composition but also alters its functional diversity. Clustering based subsystem were the most abundant category in SEED subsystems found in the polluted metagenomes (P1 and P2) and is previously reported in metagenomic studies (Delmont et al. 2012; Lavery et al. 2012). This can be attributed to the presence of many genes with unknown function due to the coupling of functional genes causing co-localization of conserved patterns in numerous genomes (Gerdes et al. 2011). However, this also proves that our understanding on the composition and function of soil microbiota is still indefinite as highlighted before in many studies (Buckley and Schmidt 2003). Upon scrutinizing the functional profile of the metagenomes for their catabolic potential, metabolism of aromatic compound with a preponderance of peripheral pathways for catabolism of aromatic compounds was observed. The genes belonging to n-phenyl alkanoic acid degradation were significantly higher and the genera *Pseudomonas* are the major group responsible for this catabolism (Olivera et al. 2001). A rise in the transcripts involved in the metabolism of aromatic compounds was observed when the soil samples were amended with phenanthrene by Menezes et al. (2012). Enrichment of genes encoding for enzymes involved in aromatic compound metabolism, stress tolerance and multidrug resistance (MDR) were distinguished with increase in salinity levels and hexachlorocyclohexane (HCH) contamination (Sangwan et al. 2012).

Ecological pressures such as thermal stress, UV radiation and pollution often cause oxidative stress, a major component of stress responses in microorganisms (Lesser 2006). As observed in this functional category of stress responses, P1, C1 and C2 were enriched with genes encoded for regulation in response to oxidative stress whilst protection from reactive oxygen species (ROS) was abundant in P2 metagenome. The enrichment of genes responsible for glutathione: non redox reactions, glutathione: redox cycle and glutaredoxins in the pristine samples signify that glutathione have significant function in ROS scavenging and act as a redox buffer to maintain the cellular balance (Jozefczak et al. 2012). Occurrence of ruberythrin encoding functional genes in P2 metagenome can be endorsed as an oxidative stress response system that includes nonheme iron proteins like rubrerythrin (Rbr) and rubredoxin oxidoreductase (Rbo) usually found in microaerophiles and anaerobes (Lumppio et al. 2000). Both the polluted metagenome depicted resistance to heavy metals (e.g. copper, zinc, cadmium, mercury and arsenic) whilst the two samples from the pristine environment showed presence of genes conferring resistance to antibiotics (e.g. fluoroquinolones and multi-drug efflux pumps). The simultaneous enrichment of oxidative stress response genes and heavy metal resistance suggests a strong correlation between responses to these two types of stresses.

Heavy metals such as lead, cadmium, mercury induce the production of ROS by reducing the other antioxidants in the cell (Ercal et al. 2001). Higher abundance of cobalt-zinc-resistance genes were observed in the polluted metagenomes which was also reported by Gillan et al. (2015) where they deciphered the adaptation of microbial community to metals. Contaminated mangrove sediment metagenomes from Brazil revealed the presence of cobalt-zinc-cadmium resistance genes (Cabral et al. 2016). A higher representation of copper homeostasis genes upon prolonged exposure to metals was highlighted in the Arctic mat study (Varin et al. 2011). These findings depict that bacteria use diverse mechanisms to adapt to anthropogenic stress and the community structures are often modulated by such selective pressures.

Antibiotic resistance genes (ARGs) are generally prevalent in the environment but along with the increasing pollution, microbes resistant to antibiotics are continuously evolving (Martinez 2009). Fluoroquinolones enter the environment through its extensive use in human medicine and for treatment of animals (Sukul and Spiteller 2007). Efflux pump related genes are usually over-expressed by bacteria as a mechanism to deal with antibiotic resistance to extrude out the toxic substrates (Webber and Piddock 2003; Alekshun and Levy 2007). Amos et al. (2014) studied the diversity of ARGs across the downstream of a river receiving the wastewater treatment plant effluents and concluded that water bodies are the most potential way of propagation of ARGs in the environment. The abundance of antibiotic resistance genes and multidrug efflux pumps in the metagenomes not only reveal the contamination through human or animal feces (Czekalski et al. 2012), but also can be due to the pharmaceutical waste discharges into the Amlakhadi canal. The functional co-relation using SEED subsystem indicates that P1 and C1 sample were highly similar in their function reflecting the similar conditions in both the sites. However, P2 and C2 had the lowest correlation which further suggests that both the conditions differ from each other. C1 and C2 also revealed some level of correlation amongst their functions. These data reveal that polluted samples are somehow closely related to control C1, possibly due to extensive long term contamination prevalent in the Amlakhadi canal that has wide spread in the nearby area from where the pristine samples were sampled within 1km range. The contaminated site being an open stream and experience the strong interference of water currents induced by the industrial effluent discharges, which brings in heavily toxic compounds that can spread across the area and imply great contamination risks for groundwaters as well (Romic and Romic 2003).

The KEGG analysis revealed nearly 40 enzymes participating in the benzoate, aminobenzoate and toluene and xylene degradation pathway that were mapped from the annotated reads of all the metagenomes. However, higher occurrence was obtained in the P1, P2 and C1 metagenome and very low mapping was observed in C2 metagenome. Many of the complex xenobiotic aromatic compounds breakdown into benzoate through various degradation pathways which then confers itself as an intermediate in the metabolism of aromatic compounds (Xu et al. 2015). As observed in Supplementary Table S4, it is expected that distinct microbes catabolize different xenobiotics or one may convert them into a lesser complex form which will be further degraded by the other microbe. This synergism allows the survival of many microorganisms that do not contribute in any degradation pathway but they maintain syntrophic associations with other microorganisms (Shah et al. 2013).

Over the past decade, it has been quite evident that enzyme sequence diversity is gaining much attention with the advances in metagenomic sequencing and earlier this functional diversity could not be captured by conventional techniques (Ufarte et al. 2015). The enzymes mapped in our study through metagenome datasets reflects the catabolic potential of the inherent microbial community residing in the polluted stream of Amlakhadi Canal (Table 1). Carboxymethylenebutenolidase being the most abundant enzyme in all the metagenomes, catalyzes the degradation of benzoate, hexachlorocyclohexane and 1,4-dichlorobenzene degradation (Ntougias et al. 2014) and has been reported to be found in the intracellular and cell wall fractions of *Burkholderia vietnamiensis* analyzed through proteomics (Wickramasekara et al. 2011). Other enzymes listed contribute in the cleavage of the intradiol ring of aromatic compounds such as catechol 1,2-dioxygenase, hydroxyquinol 1,2-dioxygenase, α-subunit), extradiol ring-cleavage enzymes such as catechol 2,3-dioxygenase, protocatechuate 4,5-dioxygenase and monooxygenases such as pentachlorophenol monooxygenase, p-hydroxybenzoate 3-monooxygenase and phenol 2-monooxygenase. Many catabolic enzymes are exploited recently for their application in the field of bioremediation (Singh et al. 2010; Nagayama et al. 2015; Santos et al. 2015; Duarte et al. 2017) and such studies further enable us to understand the role of enzymes and link their function to the ecosystem.

## 5.0 Conclusion

In conclusion, we scrutinized the microbial taxonomic and functional diversity of a long term industrially contaminated river tributary with respect to time. Prominent genera shifts were observed in the polluted samples as compared to the pristine samples upon analyzing the seasonal diversity. Functional profiles revealed the metabolic potential of the niche reflecting the catabolic resilience of the inherent microbiota. However, this study provides the information on the taxonomic identification and their functions in the polluted niche, comprehensive understanding based on the metatranscriptome and metaproteome profile will ensnare the remaining unknown functions towards environmental stress across space and time.

## Acknowledgements

This work was supported by the Department of Biotechnology (DBT) grant (BT/1/CEIB/09/V/05) from the Ministry of Science and Technology, New Delhi, India. Jenny Johnson acknowledges Department of Science and Technology (DST), New Delhi for INSPIRE fellowship.

## Author contribution statement

KRJ & DM conceived the concept of the study. JJ & KRJ performed the sampling. JJ preformed the experiments and drafted initial manuscript. JJ, KRJ, AP & NP performed the sequencing. KRJ, CJ & DM supervised the study, refined and edited the manuscript. All authors read and approved the final manuscript.

## Supplementrary Data

### 2.0 Materials and method

#### 2.6 Studying the culturable fraction of bacterial community from polluted and pristine soil samples

Soil samples (1 gm) from polluted and pristine environment were serially diluted (to obtain the colonies in the range between 30 and 300) and appropriate quantity was spreaded on different high and low nutrient media and were incubated at different temperatures (20, 37, 40 °C) for 1 to 7 days. Distinct isolated bacterial colonies with discrete morphology were screened to purity, genomic DNA was isolated and 16S rRNA gene was amplified using universal primers and sequenced as described in Desai et al. (2009).

## Supplementary Figures

**Figure S1.**
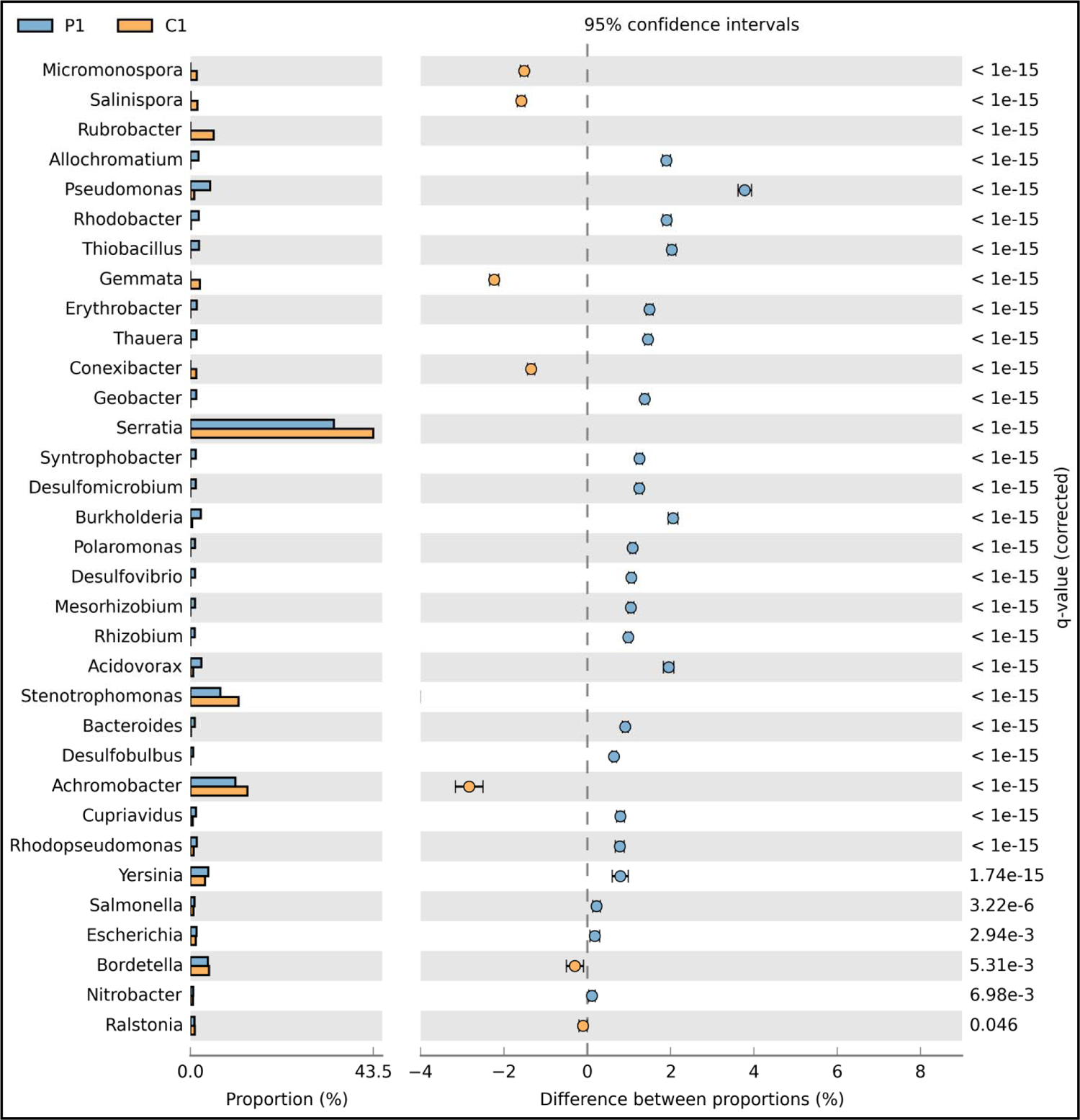
Comparative taxonomic profile of P1 (blue) and C1 (yellow) metagenomes statistically (p<0.05) analysed in STAMP at genus level.

**Figure S2.**
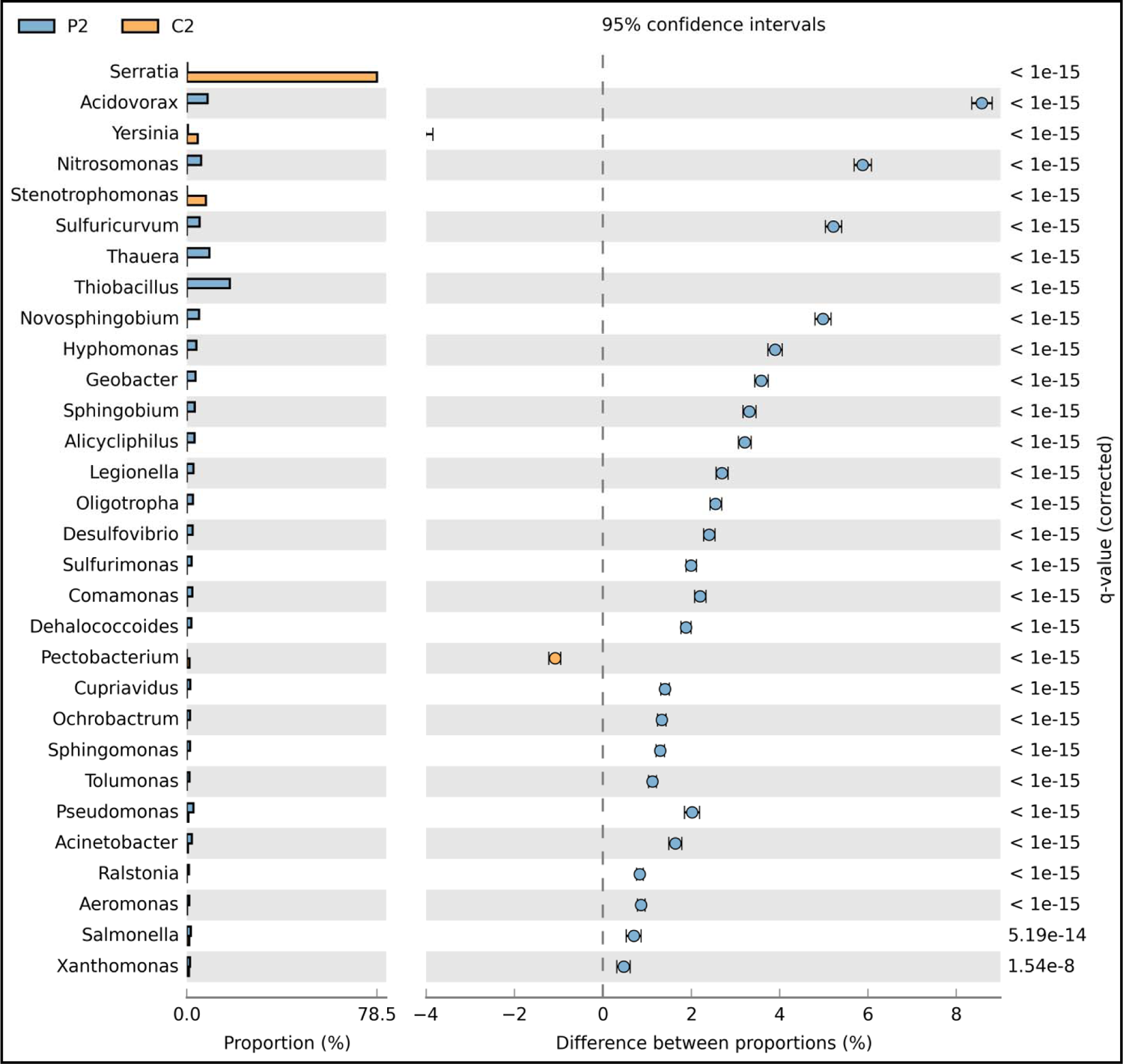
Comparative taxonomic profile of P2 (blue) and C2 (yellow) metagenomes statistically (p<0.05) analysed in STAMP at genus level.

**Figure S3.**
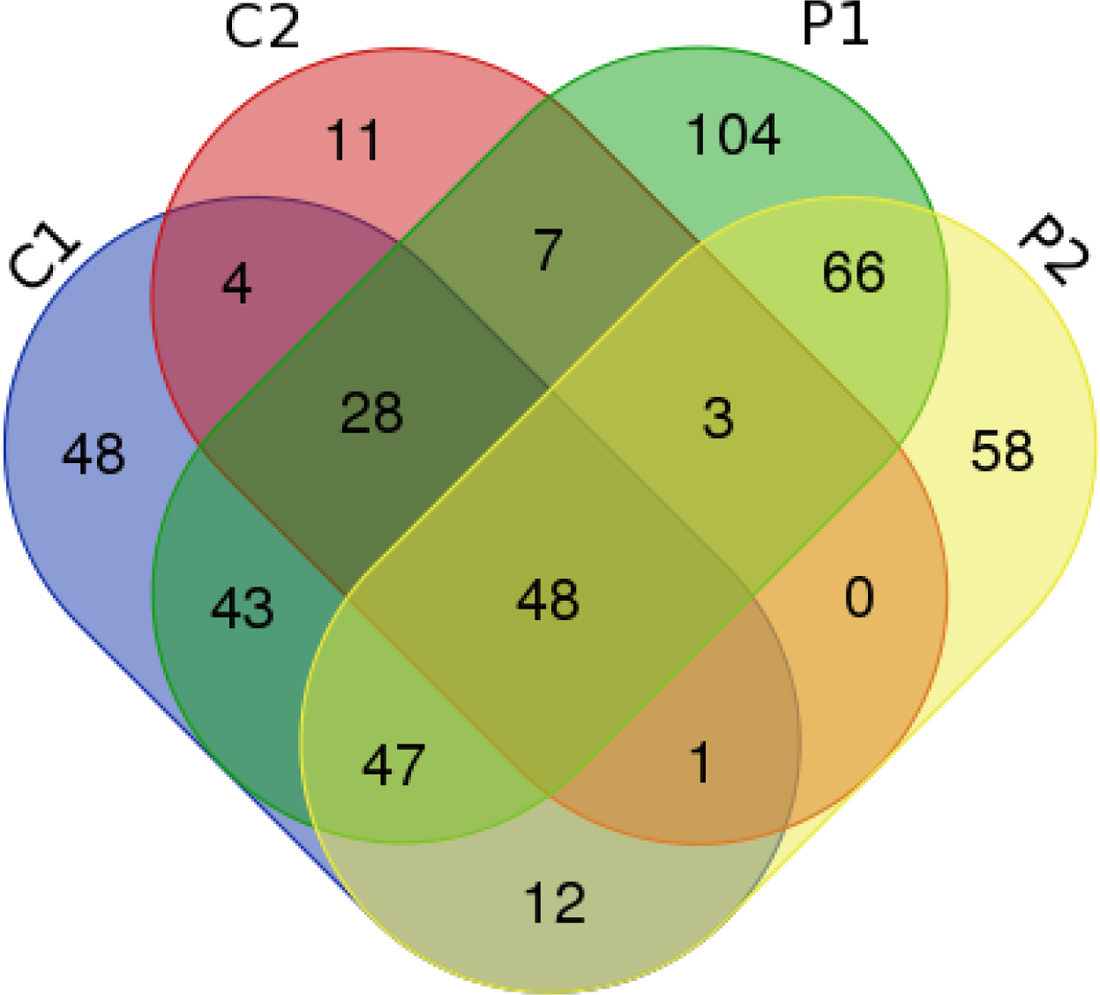
Venn diagram showing the comparison of the bacterial genera from P1, P2, C1 and C2 metagenome. All numbers in the spheres indicate the genera shared as well as unique amongst the microbial community with respect to space and time.

**Figure S4.**
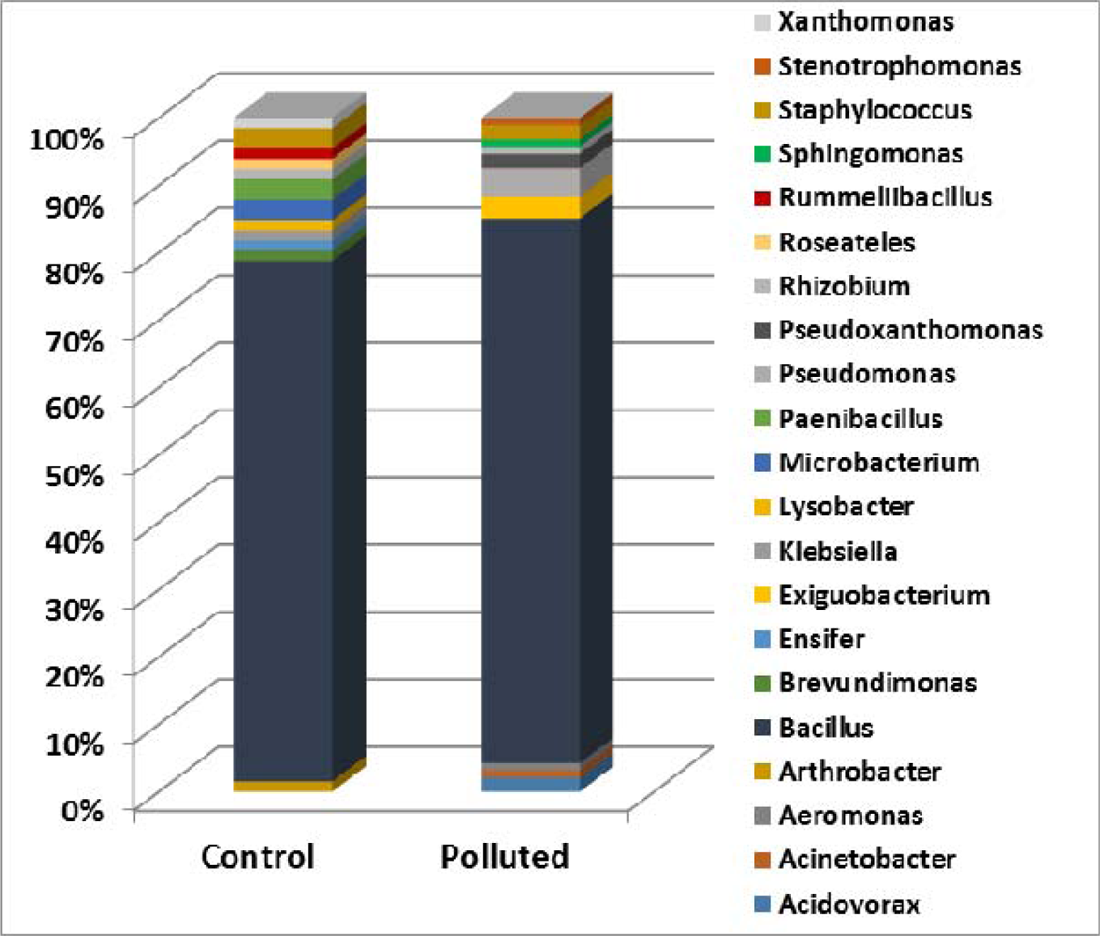
Bacterial community composition of polluted and control samples identified using culture dependent technique

**Figure S5.**
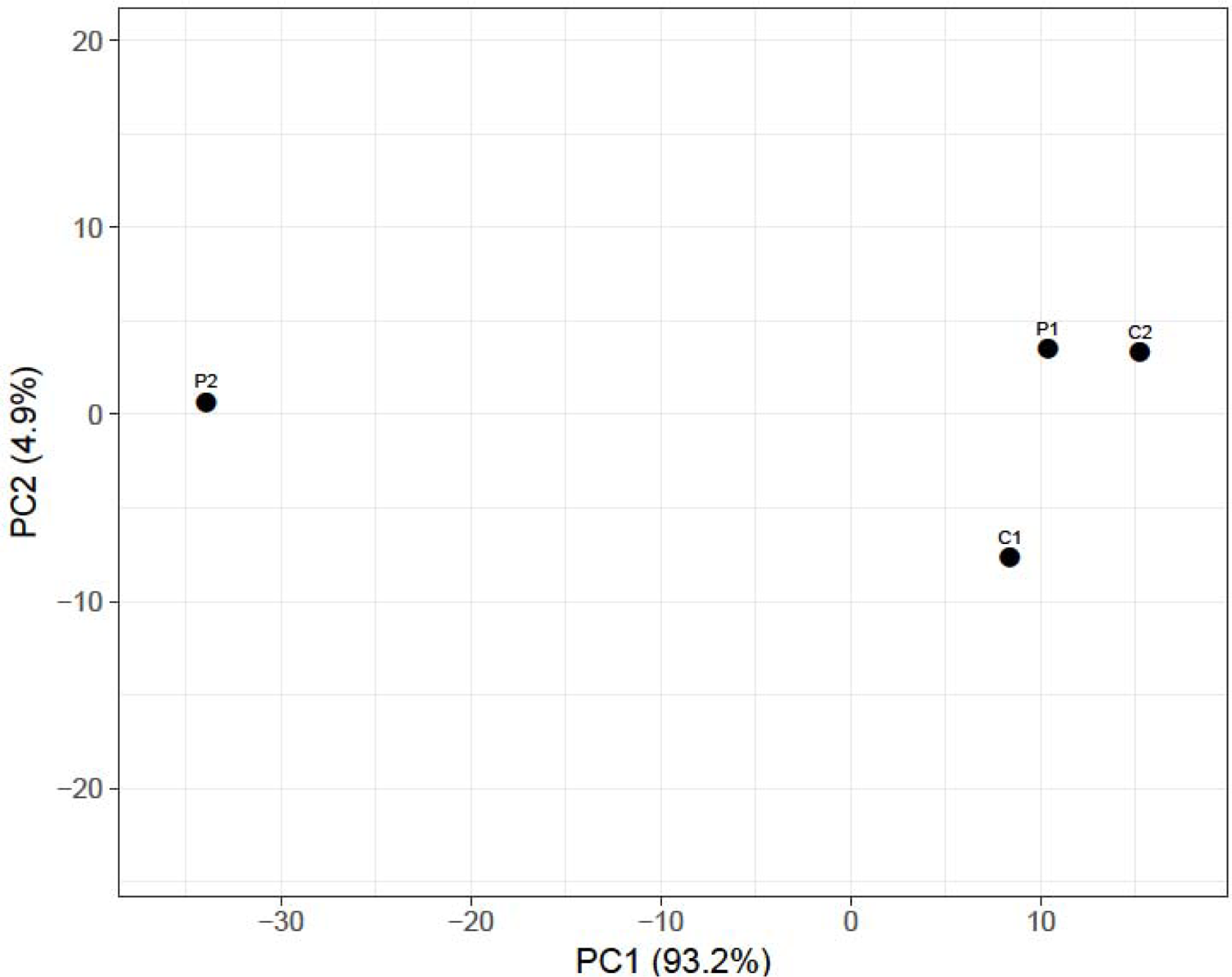
Principal component analysis (PCA) plot of the relative abundance of taxonomic composition of the metagenome analysed in PAST3 software using Bray-Curtis distance.

**Figure S6.**
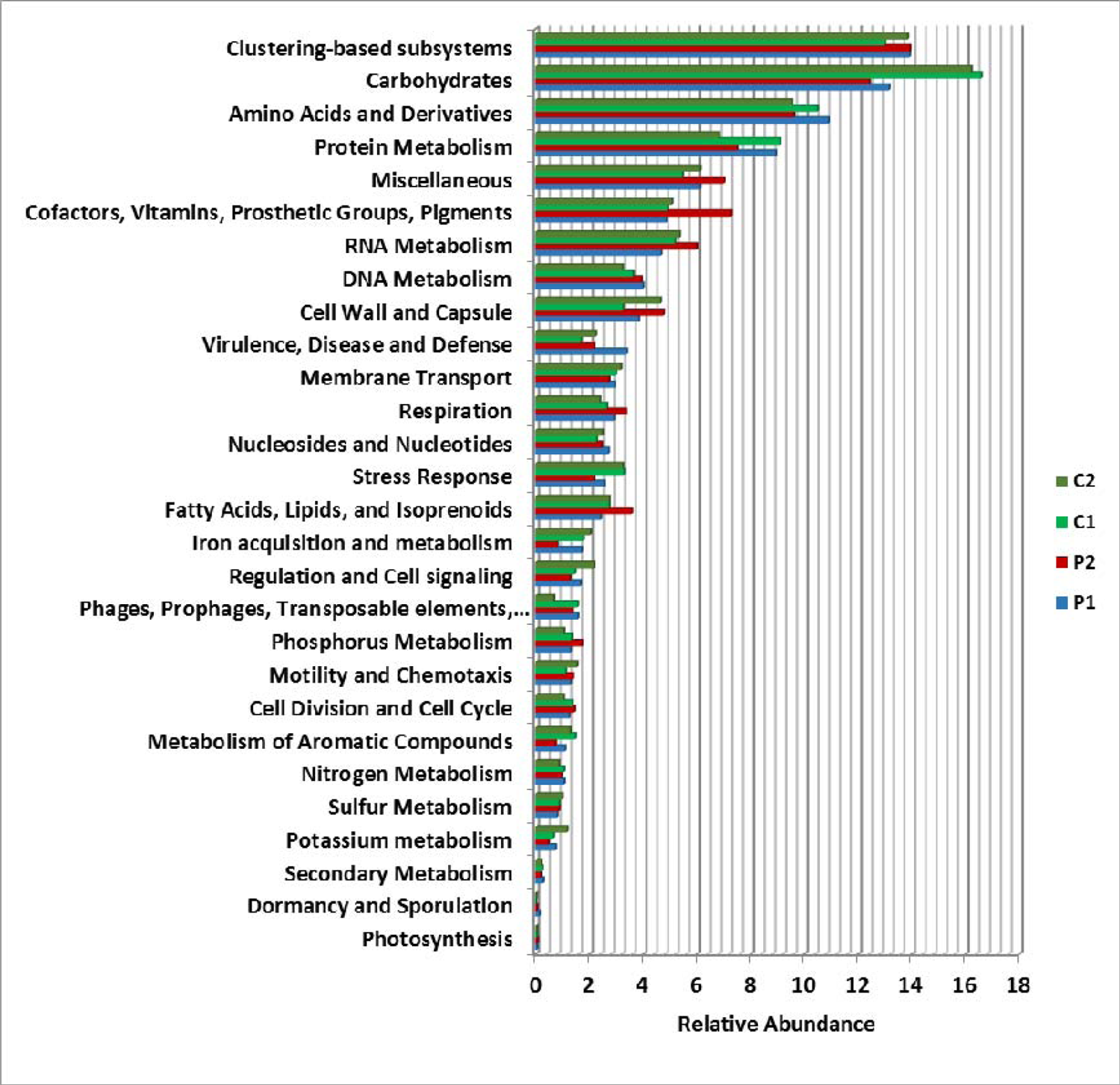
Relative abundance of functional gene categories from SEED subsystem hierarchy at Level 1 obtained for P1, P2, C1 and C2 metagenome.

**Figure S7.**
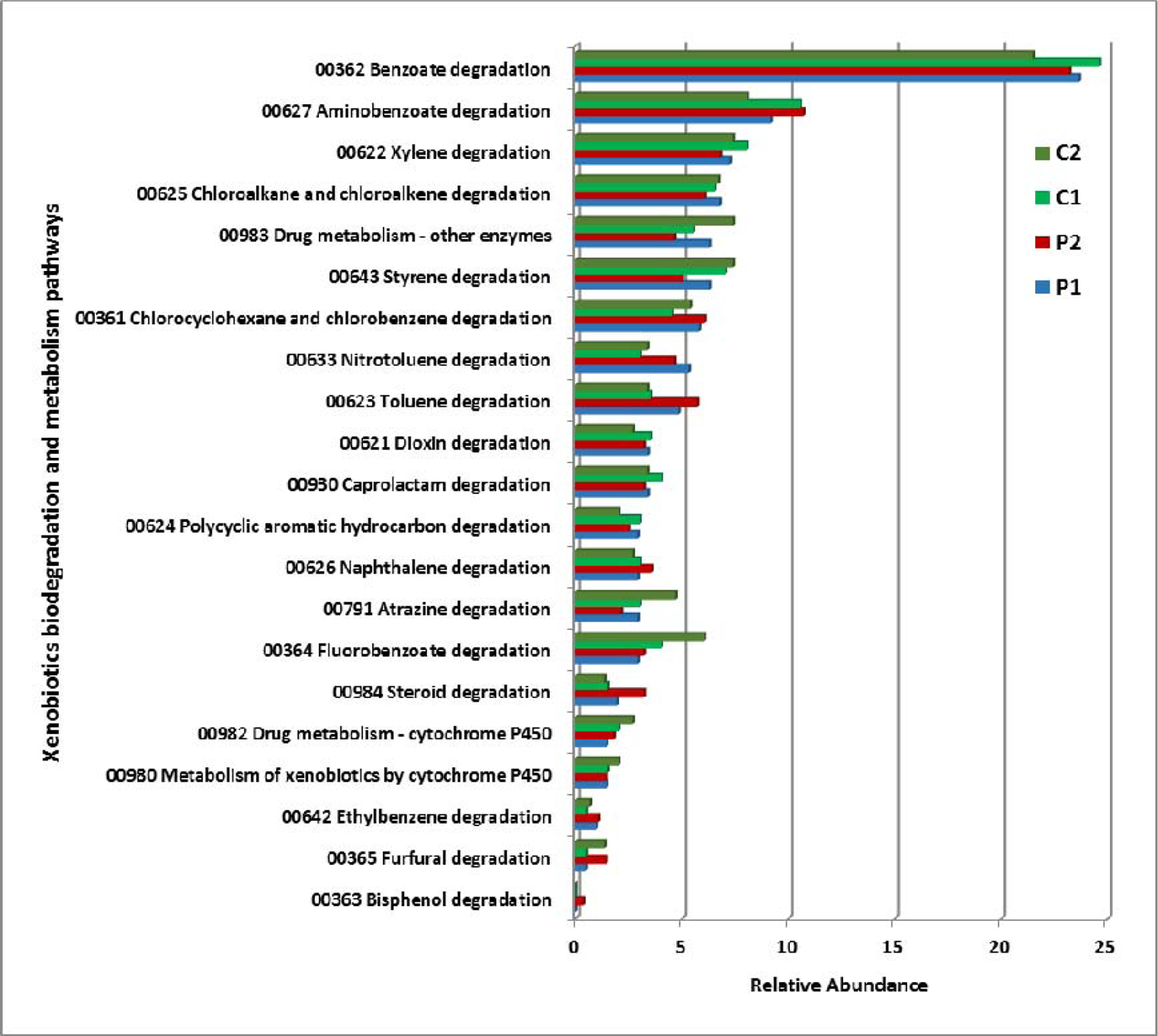
Relative abundance of pathways involved in Xenobiotics biodegradation and metabolism pathways analysed in KEGG pathway by KEGG orthology.

**Figure S8.**
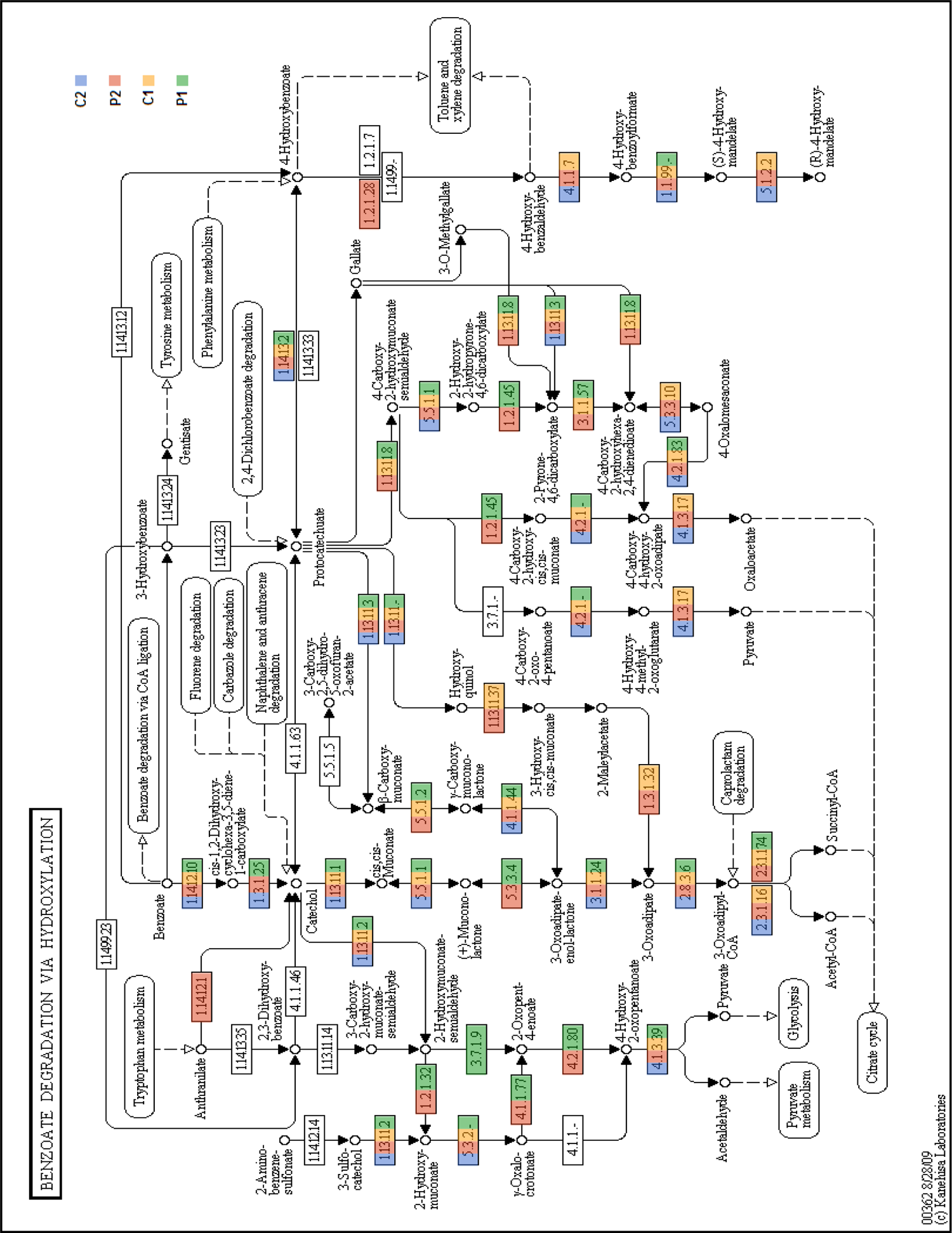
Schematic representation from KEGG pathways showing enzymes mapped for benzoate degradation via hydroxylation pathway. Mapped enzymes in each metagenome are highlighted using different colour code.

**Figure S9.**
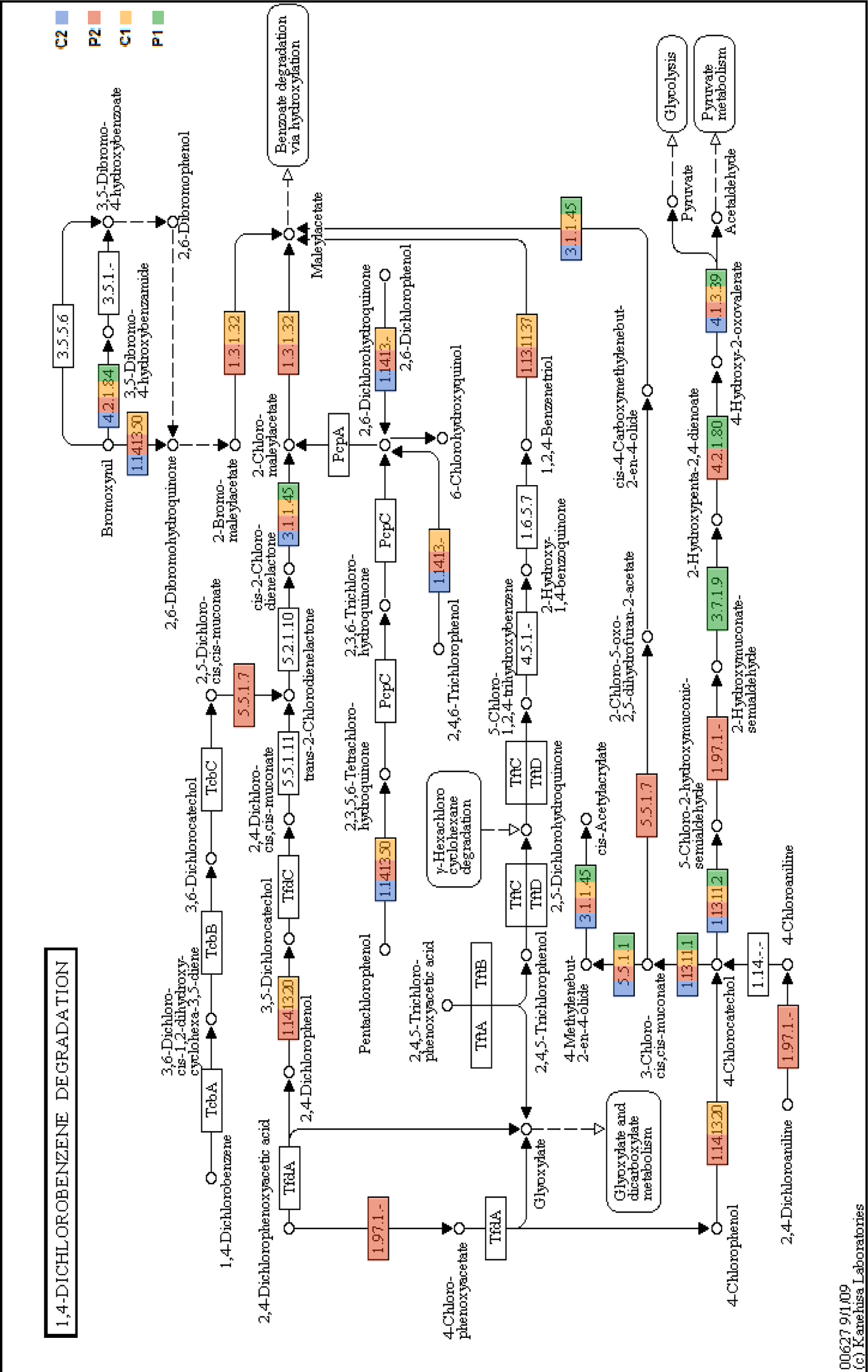
Schematic representation from KEGG pathways showing enzymes mapped for aminobenzoate degradation (1,4 Dichlorobenzene degradation). Mapped enzymes in each metagenome are highlighted using different colour code.

**Figure S10.**
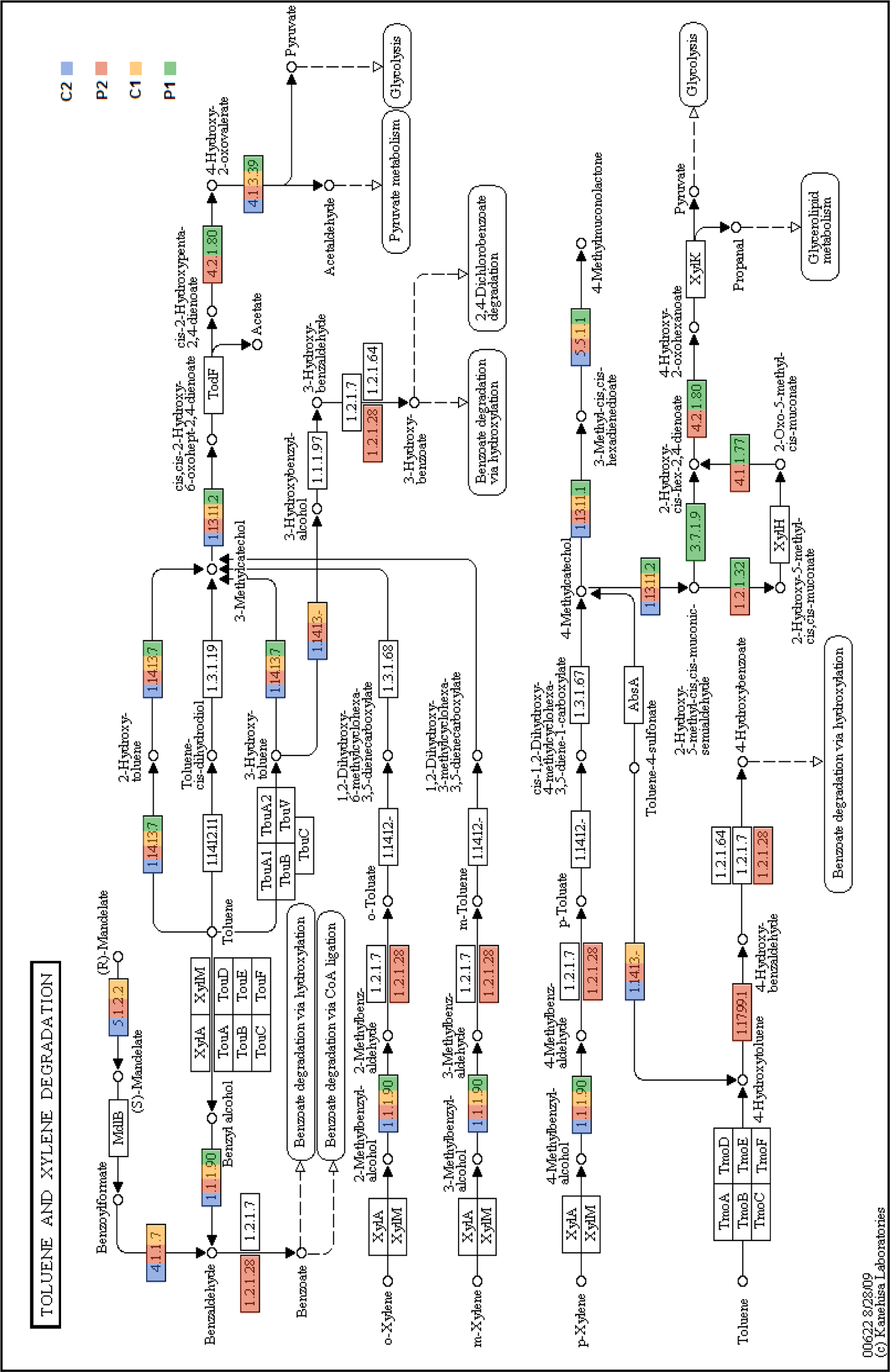
Schematic representation from KEGG pathways showing enzymes mapped for toluene and xylene degradation. Mapped enzymes in each metagenome are highlighted using different colour code.

## Supplementary Tables

**Table S1.** Physico-chemical characteristics of soil samples collected from Amlakhadi Canal

| Sample : | Amlakhadi-Control | Amlakhadi-SI | Amlakhadi-SII | Amlakhadi-SIII |
| --- | --- | --- | --- | --- |
| Variables | Physical Characteristics |  |  |  |
| pH | 7.953 | 8.173 | 8.405 | 7.877 |
| EC (ms) | 10.11 | 48.1 | 16.6 | 37.3 |
| Salinity (ppt) | 0 | 0 | 0 | 0 |
| Moisture (%) | 4.540 | 10.690 | 4.989 | 14.955 |
| WHC (%) | 50.00 | 29.00 | 55.00 | 41.00 |
| BD (g/cm<sup>3</sup>) | 1.369 | 1.333 | 1.190 | 1.538 |
| SG | 1.779 | 3.063 | 1.931 | 1.668 |
|  | Chemical Characteristics |  |  |  |
| TOC (%) | 0.66 | 1.35 | 0.88 | 1.09 |
| Na (g/kg) | 11.57 | 15.55 | 11.025 | 11.65 |
| K (g/kg) | 0.00025 | 0.0005 | 0.03 | 0.05 |
| Ca (g/kg) | 9.25 | 0.375 | 20.65 | 2.19 |
| P (g/kg) | 0.002 | 0.001 | 0.004 | 0.003 |
| S (g/kg) | 0.013 | 0.015 | 0.011 | 0.010 |
| Cl (g/kg) | 0.031 | 0.067 | 0.063 | 0.071 |
|  | Heavy metals |  |  |  |
| Cr (g/kg) | 0.821 | 0.939 | 0.896 | 0.825 |
| Co (g/kg) | 0.589 | 0.568 | 0.566 | 0.569 |
| Fe (g/kg) | 0.027 | 0.092 | 0.832 | 0.627 |
| Cu (g/kg) | 0.63 | 2.096 | 0.625 | 1.127 |
| Mn (g/kg) | 2.212 | 1.365 | 2.512 | 1.495 |
| Zn (g/kg) | 0.343 | 1.421 | 0.39 | 0.678 |
| Cd (g/kg) | BDL | BDL | BDL | BDL |
| SI - Site 1 | EC = | Electrical Conductivity | SG = | Specific gravity |
| SII - Site 2 | WHC = | Water Holding Capacity | TOC = | Total Organic Carbon |
| SIII - Site 3 | BD = | Bulk Density | BDL = | Below Detectable Level |

**Table S2.**
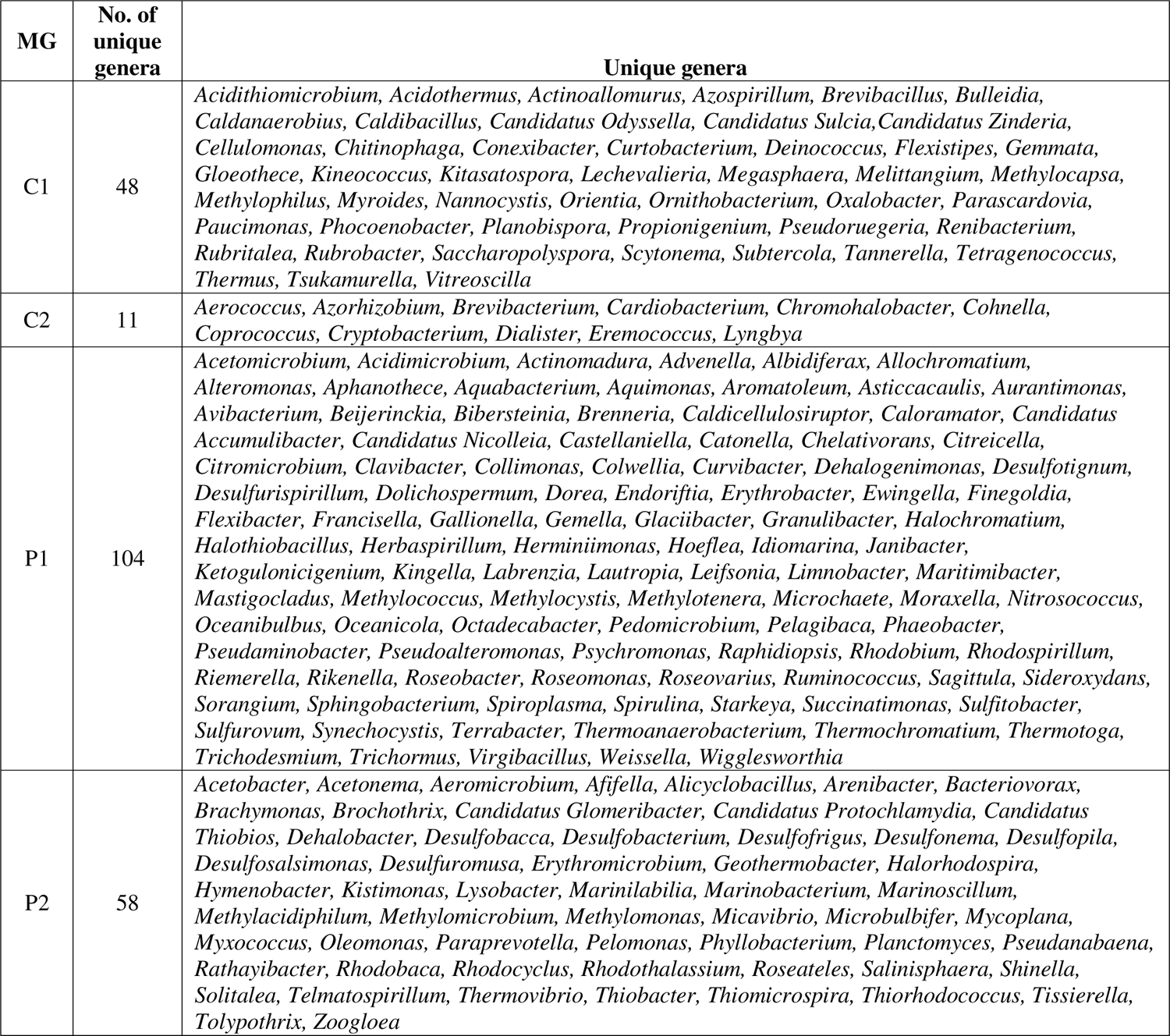
List of unique genera in the polluted and control metagenomes of both the seasons

| MG | No. of unique genera | Unique genera |
| --- | --- | --- |
| C1 | 48 | *Acidithiomicrobium*, *Acidothermus*, *Actinoallomurus*, *Azospirillum*, *Brevibacillus*, *Bulleidia*, *Caldanaerobius*, *Caldibacillus*, *Candidatus Odysella*, *Candidatus Sulcia*, *Candidatus Zinderia*, *Cellulomonas*, *Chitinophaga*, *Conexibacter*, *Curtobacterium*, *Deinococcus*, *Flexistipes*, *Gemmata*, *Gloeotheca*, *Kineococcus*, *Kitasatospora*, *Lechevalieria*, *Megasphaera*, *Melittangium*, *Methylocapsa*, *Methylophilus*, *Myroides*, *Nannocystis*, *Orientia*, *Ornithobacterium*, *Oxalobacter*, *Parascardovia*, *Paucimonas*, *Phocoenobacter*, *Planobispora*, *Propionigenium*, *Pseudoruegeria*, *Renibacterium*, *Rubritalea*, *Rubrobacter*, *Saccharopolyspora*, *Scytonema*, *Subtercola*, *Tannerella*, *Tetragenococcus*, *Thermus*, *Tsukamurella*, *Vitreoscilla* |
| C2 | 11 | *Aerococcus*, *Azorhizobium*, *Brevibacterium*, *Cardiobacterium*, *Chromohalobacter*, *Cohnella*, *Coprococcus*, *Cryptobacterium*, *Dialister*, *Eremococcus*, *Lyngbya* |
| P1 | 104 | *Acetomicrobium*, *Acidimicrobium*, *Actinomadura*, *Advenella*, *Albidiferax*, *Allochromatium*, *Alteromonas*, *Aphanotheca*, *Aquabacterium*, *Aquimonas*, *Aromatoleum*, *Asticcacaulis*, *Aurantimonas*, *Avibacterium*, *Beijerinckia*, *Bibersteinia*, *Brenneria*, *Caldicellulosiruptor*, *Caloramator*, *Candidatus Accumulibacter*, *Candidatus Nicolleia*, *Castellaniella*, *Catonella*, *Chelativorans*, *Citricella*, *Citromicrobium*, *Clavibacter*, *Collimonas*, *Colwellia*, *Curvibacter*, *Dehalogenimonas*, *Desulfotignum*, *Desulfurispirillum*, *Dolichospermum*, *Dorea*, *Endoriftia*, *Erythrobacter*, *Ewingella*, *Finegoldia*, *Flexibacter*, *Francisella*, *Gallionella*, *Gemella*, *Glaciibacter*, *Granulibacter*, *Halochromatium*, *Halothiobacillus*, *Herbaspirillum*, *Herminiimonas*, *Hoeflea*, *Idiomarina*, *Janibacter*, *Ketogulonicigenium*, *Kingella*, *Labrenzia*, *Lautropia*, *Leifsonia*, *Limnobacter*, *Maritimibacter*, *Mastigocladus*, *Methylococcus*, *Methylocystis*, *Methylotenera*, *Microchaete*, *Moraxella*, *Nitrosococcus*, *Oceanibulbus*, *Oceanicola*, *Octadecabacter*, *Pedomicrobium*, *Pelagibaca*, *Phaeobacter*, *Pseudaminobacter*, *Pseudoalteromonas*, *Psychromonas*, *Raphidiopsis*, *Rhodobium*, *Rhodospirillum*, *Riemerella*, *Rikenella*, *Roseobacter*, *Roseomonas*, *Roseovarius*, *Ruminococcus*, *Sagittula*, *Sideroxydans*, *Sorangium*, *Sphingobacterium*, *Spiroplasma*, *Spirulina*, *Starkeya*, *Succinatimonas*, *Sulfitobacter*, *Sulfurovum*, *Synechocystis*, *Terrabacter*, *Thermoanaerobacterium*, *Thermochromatium*, *Thermotoga*, *Trichodesmium*, *Trichormus*, *Virgibacillus*, *Weissella*, *Wigglesworthia* |
| P2 | 58 | *Acetobacter*, *Acetonema*, *Aeromicrobium*, *Ajifella*, *Alicyclobacillus*, *Arenibacter*, *Bacteriovorax*, *Brachymonas*, *Brochothrix*, *Candidatus Glomeribacter*, *Candidatus Protochlamydia*, *Candidatus Thiobios*, *Dehalobacter*, *Desulfobacca*, *Desulfobacterium*, *Desulfofrigus*, *Desulfonema*, *Desulfopila*, *Desulfosalsimonas*, *Desulfuromusa*, *Erythromicrobium*, *Geothermobacter*, *Halorhodospira*, *Hymenobacter*, *Kistimonas*, *Lysobacter*, *Marinilabilia*, *Marinobacterium*, *Marinoscillum*, *Methylacidiphilum*, *Methylomicrobium*, *Methylomonas*, *Micavibrio*, *Microbulbifer*, *Mycoplana*, *Myxococcus*, *Oleomonas*, *Paraprevotella*, *Pelomonas*, *Phyllobacterium*, *Planctomyces*, *Pseudanabaena*, *Rathayibacter*, *Rhodobaca*, *Rhodocyclus*, *Rhodothalassium*, *Roseateles*, *Salinisphaera*, *Shinella*, *Solitalea*, *Telmatospirillum*, *Thermovibrio*, *Thiobacter*, *Thiomicrospira*, *Thiorhodococcus*, *Tissierella*, *Tolypothrix*, *Zoogloea* |

**Table S3.** Diversity indices of all the metagenomes determined by PAST3 software

| **Diversity Indices** | **Metagenome** |  |  |  |
| --- | --- | --- | --- | --- |
|  | P1 | C1 | P2 | C2 |
| Dominance_D | 0.06608 | 0.1684 | 0.05801 | 0.5417 |
| Simpson_1-D | 0.9339 | 0.8316 | 0.942 | 0.4583 |
| Shannon_H | 4.038 | 2.792 | 3.381 | 1.352 |
| Fisher_alpha | 43.74 | 29.05 | 31.01 | 13.72 |
| Chao-1 | 428.6 | 286.7 | 336.3 | 135.1 |

**Table S4.**
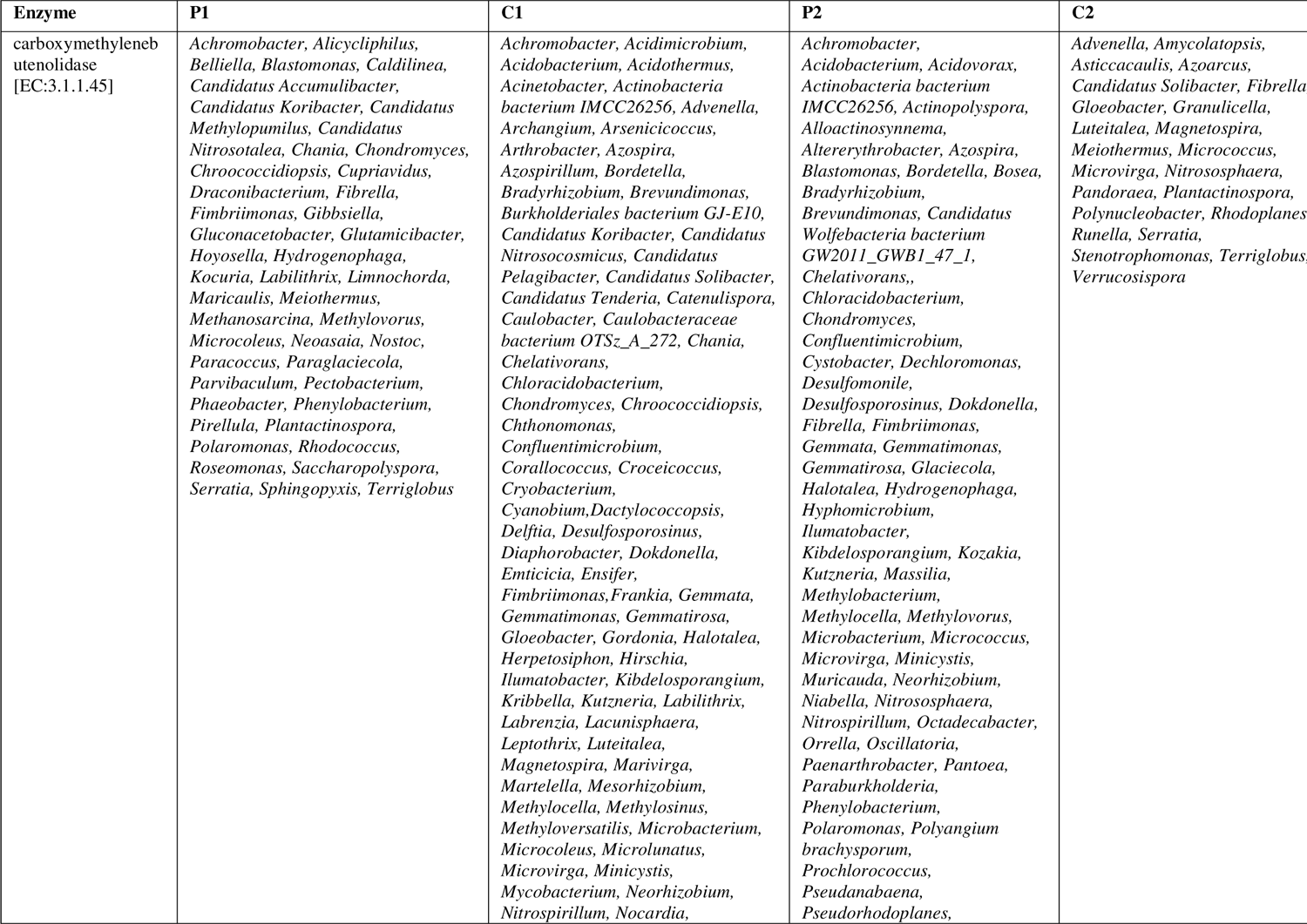
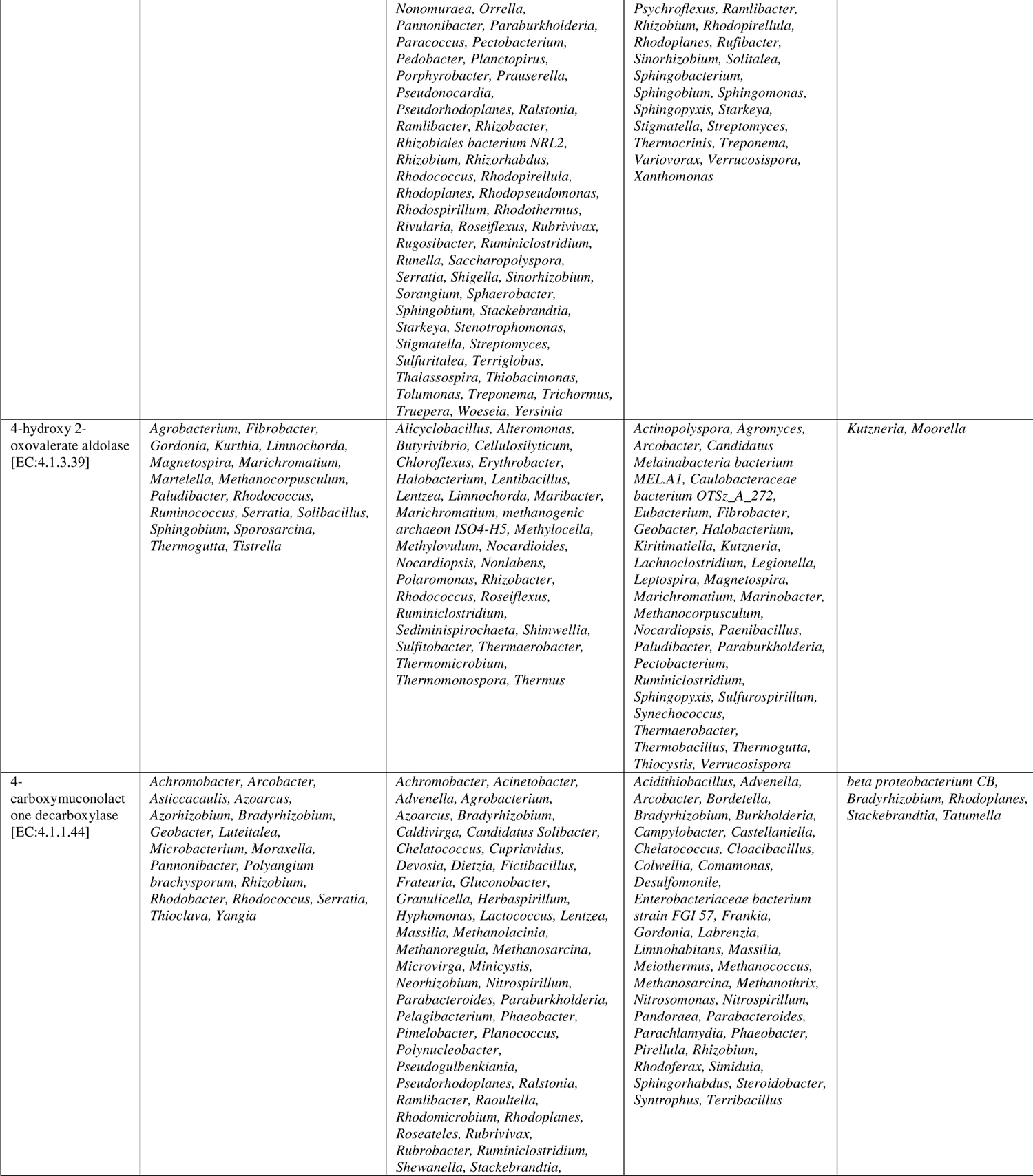
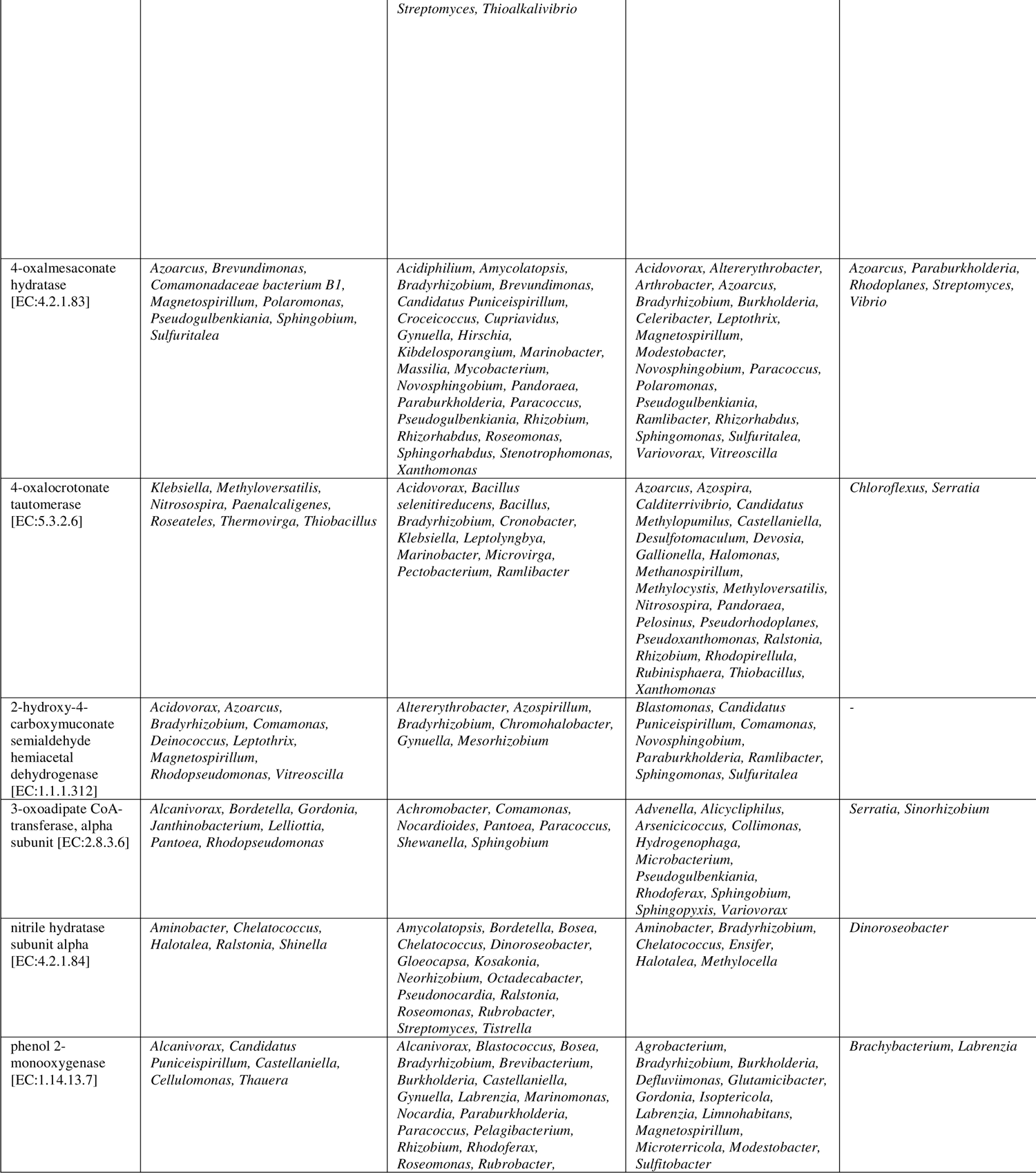
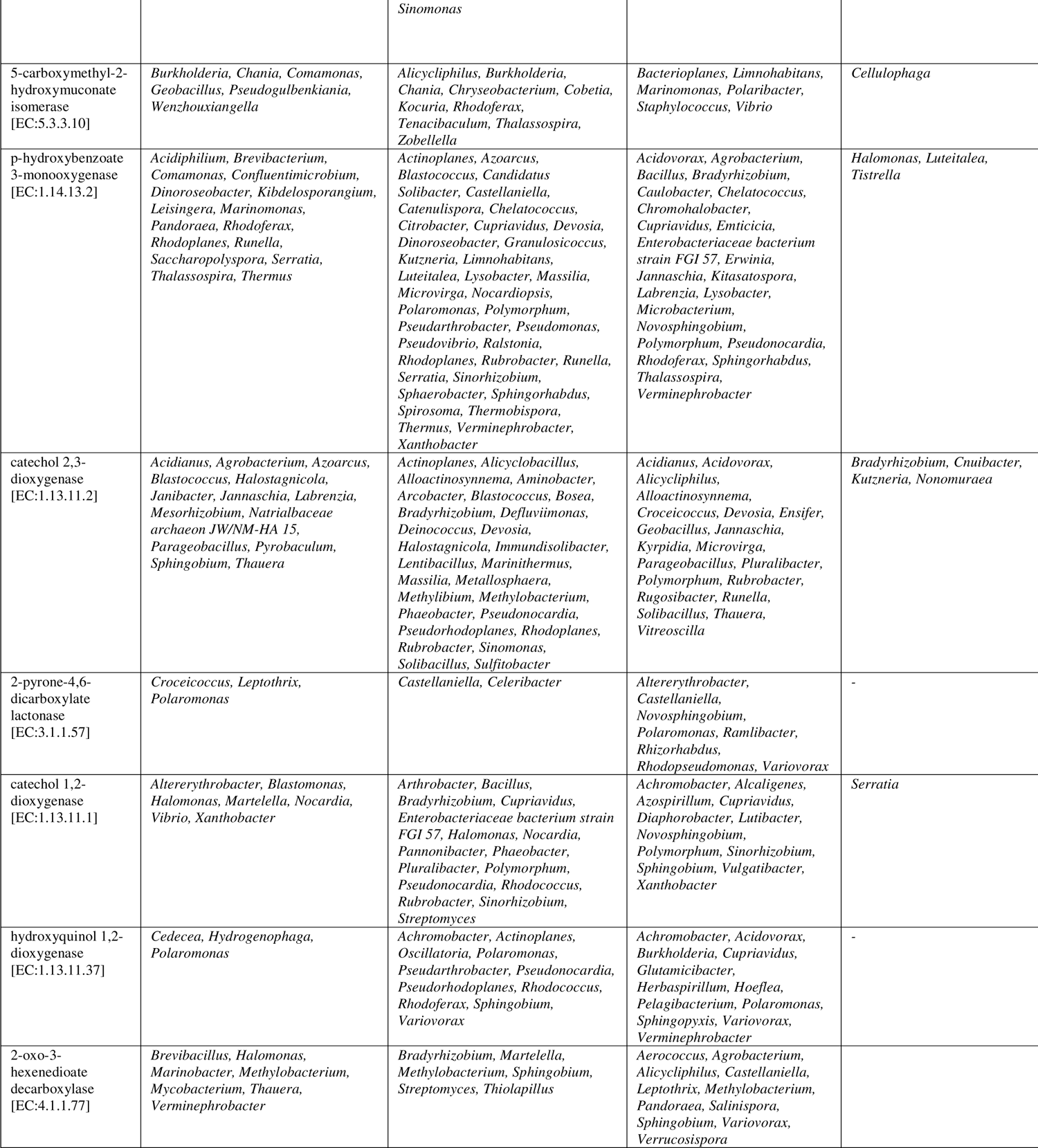
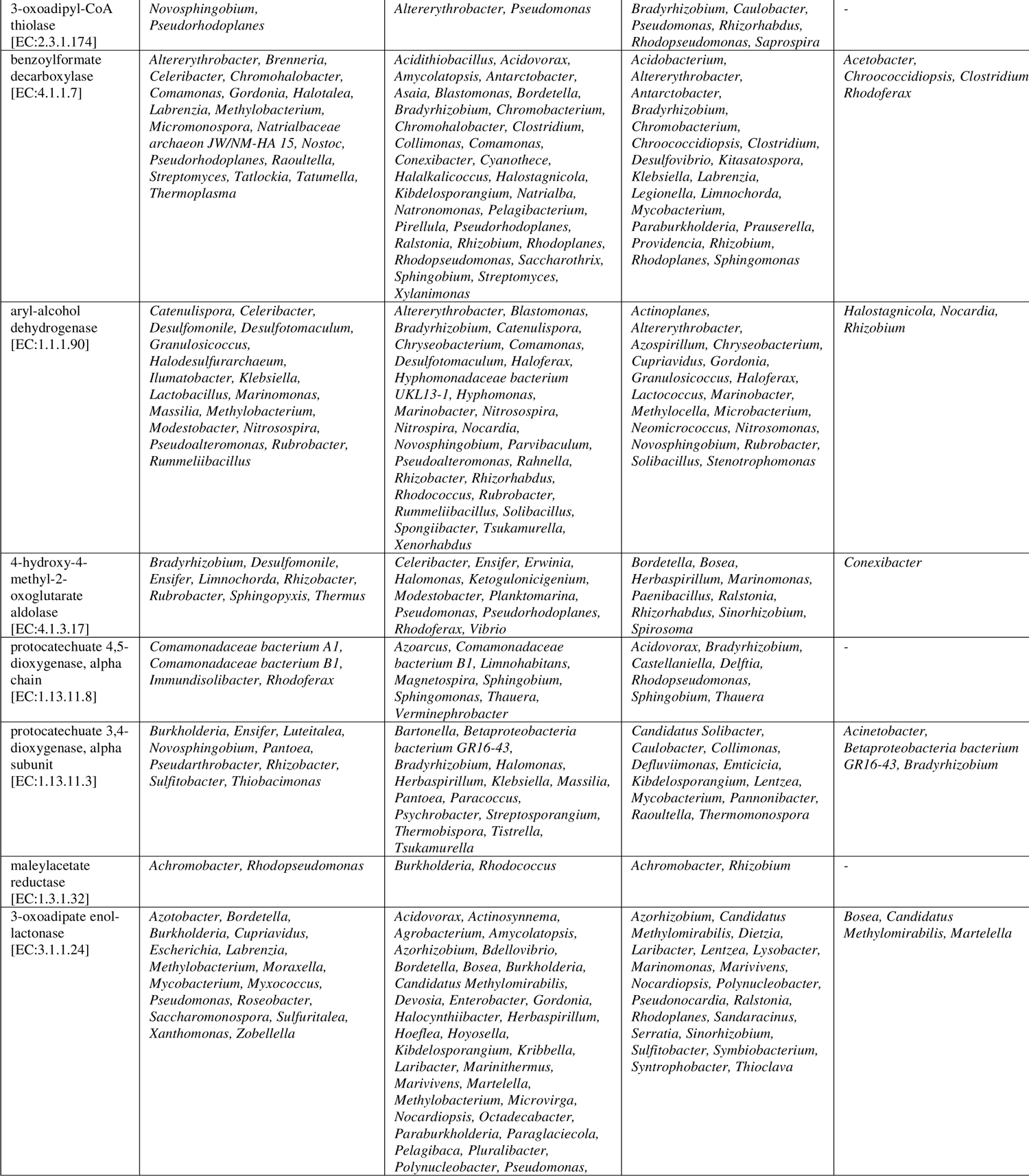
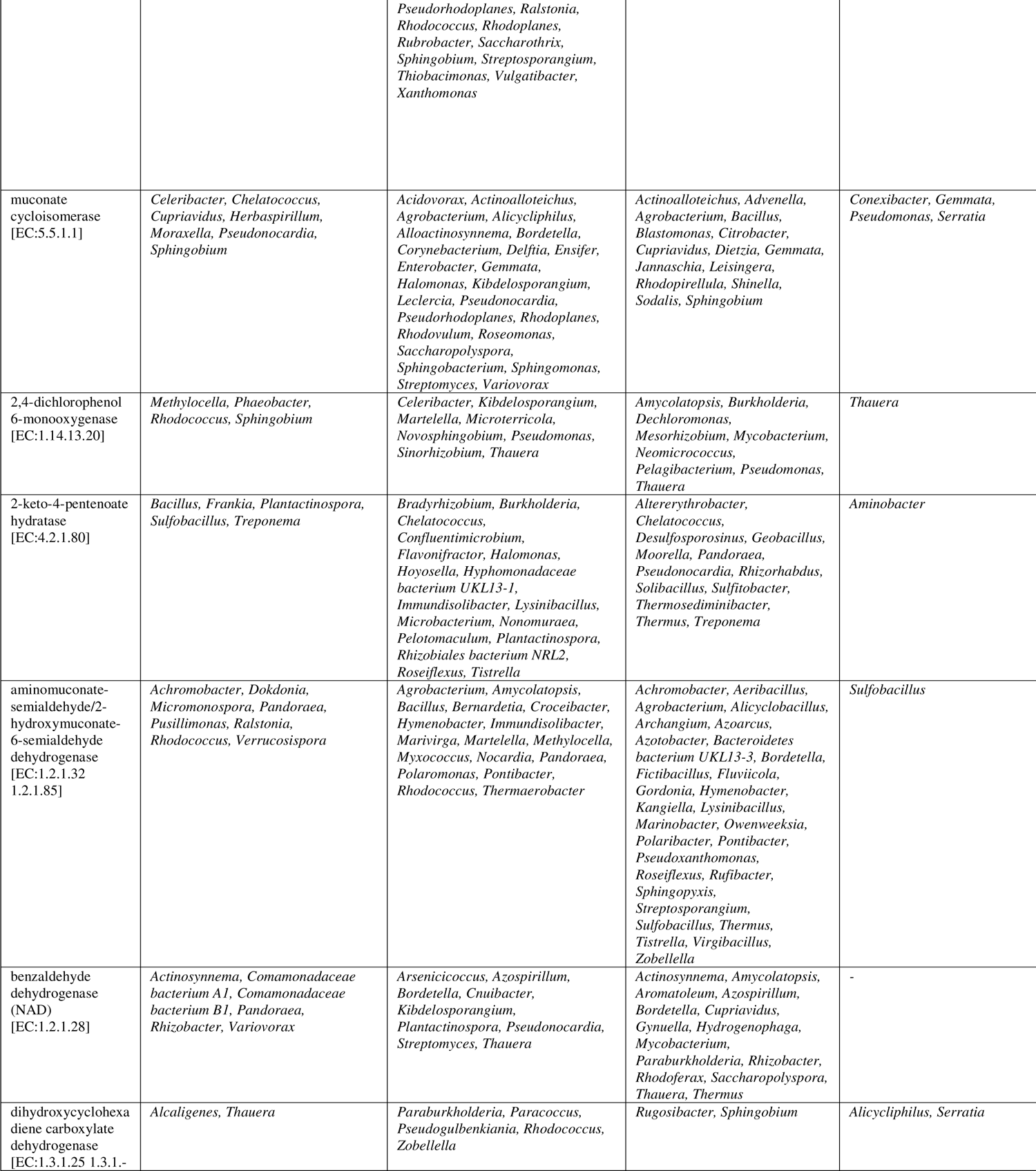
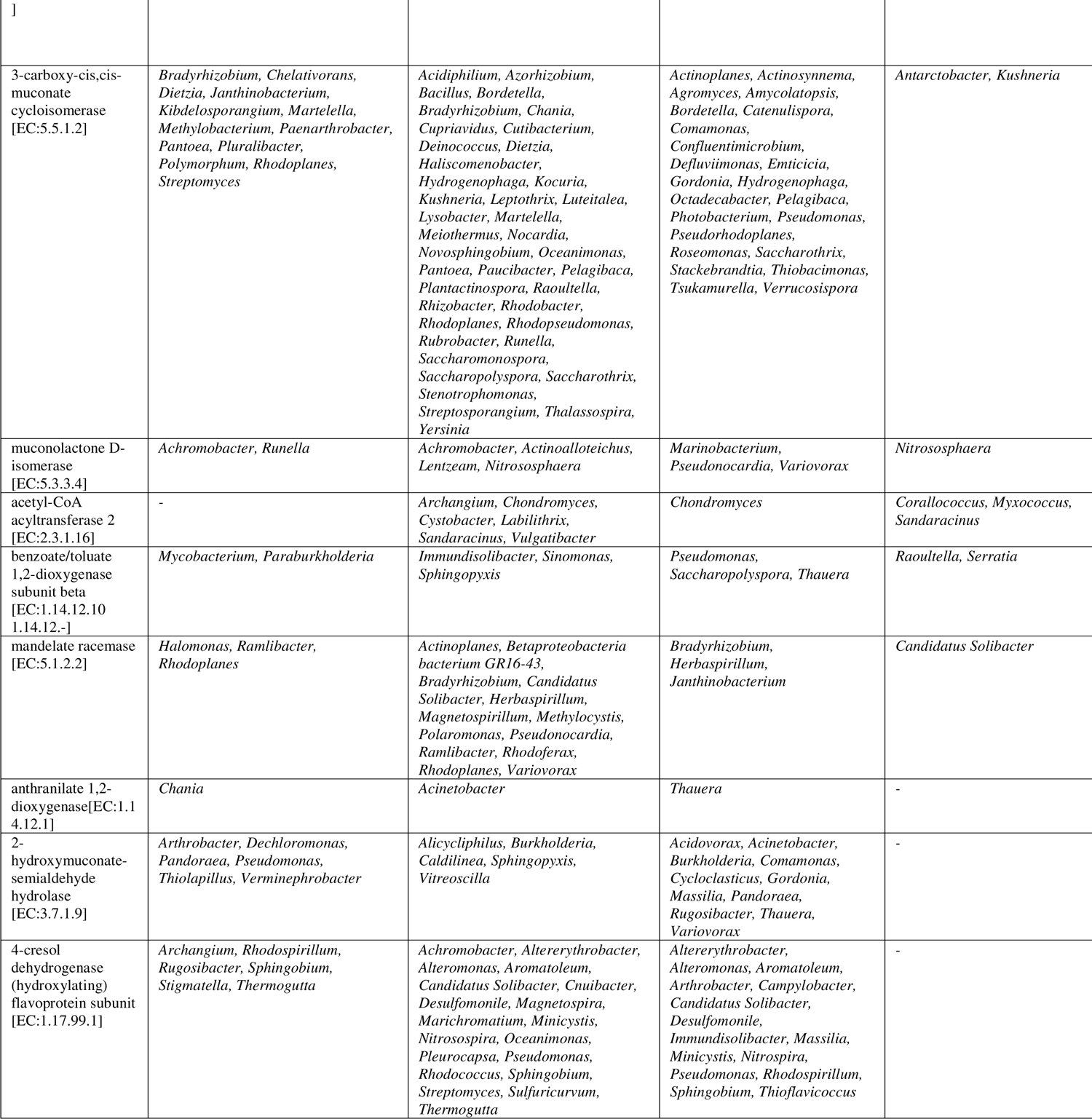
List of genera associated with the enzymes mapped for degradation of benzoate, aminobenzoate, toluene and xylene obtained from GHOSTKOALA related to KEGG pathway analysis

## References

1. Adetutu EM, Smith RJ, Weber J, Aleer, S, Mitchell JG, Ball AS, Juhasz, AL (2013) A polyphasic approach for assessing the suitability of bioremediation for the treatment of hydrocarbon-impacted soil. Science of the total environment 450:51–58.

2. Alcalde M, Ferrer M, Plou FJ, Ballesteros A (2006) Environmental biocatalysis: from remediation with enzymes to novel green processes. TRENDS in Biotechnology 24(6), 281–287.

3. Alekshun MN, Levy SB (2007) Molecular mechanisms of antibacterial multidrug resistance. Cell 128(6), 1037–1050.

4. Amos GC, Zhang L, Hawkey PM, Gaze WH, Wellington EM (2014). Functional metagenomic analysis reveals rivers are a reservoir for diverse antibiotic resistance genes. Veterinary microbiology 171(3), 441–447.

5. Aylward FO, McDonald BR, Adams SM, Valenzuela A, Schmidt, RA, Goodwin LA, Woyke, T, Currie, CR, Suen G, Poulsen M (2013). Comparison of 26 sphingomonad genomes reveals diverse environmental adaptations and biodegradative capabilities. Applied and environmental microbiology 79(12), 3724–3733.

6. Badalamenti JP, Krajmalnik-Brown R, Torres CI, Bond DR, (2015). Genomes of *Geoalkalibacter ferrihydriticus* Z-0531T and *Geoalkalibacter subterraneus* Red1T, two haloalkaliphilic metal-reducing Deltaproteobacteria. Genome announcements 3(2), e00039–15.

7. Baraniecki CA, Aislabie J, Foght JM (2002). Characterization of Sphingomonas sp. Ant 17, an aromatic hydrocarbon-degrading bacterium isolated from Antarctic soil. Microbial Ecology 43(1), 44-54.

8. Bell TH, Yergeau E, Martineau C, Juck D, Whyte LG, Greer CW (2011). Identification of nitrogen-incorporating bacteria in petroleum-contaminated arctic soils by using [15N] DNA-based stable isotope probing and pyrosequencing. Applied and environmental microbiology 77(12), 4163–4171.

9. Buckley DH, Schmidt TM (2003). Diversity and dynamics of microbial communities in soils from agro-ecosystems. Environmental Microbiology 5(6), 441–52.

10. Cabral L, Júnior GV, de Sousa, ST, Dias AC, Cadete LL, Andreote FD, Hess M, de Oliveira VM (2016). Anthropogenic impact on mangrove sediments triggers differential responses in the heavy metals and antibiotic resistomes of microbial communities. Environmental Pollution 216, 460–469.

11. Chougule AS, Jadhav SB, Jadhav JP (2014) Microbial degradation and detoxification of synthetic dye mixture by Pseudomonas sp. SUK 1. Proceedings of the National Academy of Sciences, India Section B Biological Sciences 84(4), 1059-1068.

12. CPCB (2010). Annual Water Quality Statistics of India, Central Pollution Control Board, Government of India.

13. Czekalski N, Berthold T, Caucci, S, Egli A, Bürgmann H (2012) Increased levels of multiresistant bacteria and resistance genes after wastewater treatment and their dissemination into Lake Geneva, Switzerland. Frontiers in Microbiology 3 https://doi.org/10.3389/fmicb.2012.00106

14. de Menezes A, Clipson N, Doyle E (2012) Comparative metatranscriptomics reveals widespread community responses during phenanthrene degradation in soil. Environmental microbiology 14(9), 2577–2588.

15. Delmont TO, Prestat E, Keegan KP, Faubladier M, Robe P, Clark IM, Pelletier E, Hirsch PR, Meyer F, Gilbert JA, Le Paslier, D (2012) Structure, fluctuation and magnitude of a natural grassland soil metagenome. The ISME journal 6(9), 1677–1687.

16. Desai C, Jain K, Patel B, Madamwar D (2009) Efficacy of bacterial consortium-AIE2 for contemporaneous Cr (VI) and azo dye bioremediation in batch and continuous bioreactor systems, monitoring steady-state bacterial dynamics using qPCR assays. Biodegradation 20(6), 813.

17. Desai C, Madamwar D (2007) Extraction of inhibitor-free metagenomic DNA from polluted sediments, compatible with molecular diversity analysis using adsorption and ion-exchange treatments. Bioresource Technology 98(4), 761–768.

18. Desai C, Pathak H, Madamwar D (2010). Advances in molecular and “-omics” technologies to gauge microbial communities and bioremediation at xenobiotic/anthropogen contaminated sites. Bioresource Technology 101(6), 1558–1569.

19. dos Santos DF, Istvan P, Noronha EF, Quirino BF, Krüger RH (2015) New dioxygenase from metagenomic library from Brazilian soil: insights into antibiotic resistance and bioremediation. Biotechnology letters 37(9), 1809–1817.

20. Duarte M, Nielsen A, Camarinha-Silva, A, Vilchez-Vargas R, Bruls T, Wos-Oxley ML, Jauregui R, Pieper DH (2017). Functional soil metagenomics: elucidation of polycyclic aromatic hydrocarbon degradation potential following 12 years of in situ bioremediation. Environmental Microbiology Apr 12.

21. Ercal N, Gurer-Orhan H, Aykin-Burns N (2001) Toxic metals and oxidative stress part I: mechanisms involved in metal-induced oxidative damage. Current topics in medicinal chemistry 1(6), 529–539.

22. Fierer N, Breitbart M, Nulton J, Salamon P, Lozupone C, Jones, R, Robeson M, Edwards, RA, Felts B, Rayhawk S, Knight, R (2007). Metagenomic and small-subunit rRNA analyses reveal the genetic diversity of bacteria, archaea, fungi, and viruses in soil. Applied and environmental microbiology 73(21), 7059–7066.

23. Fortin D, Davis B, Beveridge TJ (1996) Role of *Thiobacillus* and sulfate-reducing bacteria in iron biocycling in oxic and acidic mine tailings. FEMS Microbiology Ecology 21(1), 11–24.

24. Garg SK, Tripathi M, Lal N (2015). Response surface methodology for optimization of process variable for reactive orange 4 dye discoloration by *Pseudomonas putida* SKG-1 strain and bioreactor trial for its possible use in large-scale bioremediation. Desalination and Water Treatment 54(11), 3122–3133.

25. Gaston KJ (2010) Valuing common species. Science 327(5962), 154–155.

26. Gerdes S, El Yacoubi B, Bailly M, Blaby IK, Blaby-Haas, C.E., Jeanguenin, L., Lara-Núñez, A., Pribat, A., Waller, J.C., Wilke, A., Overbeek, R., 2011. Synergistic use of plant-prokaryote comparative genomics for functional annotations. BMC genomics 12(1), S2.

27. Gianoulis, TA, Raes, J, Patel, PV, Bjornson R, Korbel JO, Letunic I, Yamada T, Paccanaro A, Jensen LJ, Snyder M, Bork P (2009) Quantifying environmental adaptation of metabolic pathways in metagenomics. Proceedings of the National Academy of Sciences 106(5), 1374–1379.

28. Gillan DC, Roosa S, Kunath B, Billon G, Wattiez R (2015) The long-term adaptation of bacterial communities in metal-contaminated sediments: a metaproteogenomic study. Environmental microbiology 17(6), 1991–2005.

29. Goł biewski M, Deja-Sikora E, Cichosz M, Tretyn A, Wróbel B (2014) 16S rDNA pyrosequencing analysis of bacterial community in heavy metals polluted soils. Microbial ecology 67(3), 635–647.

30. Gomez-Alvarez V, Teal TK, Schmidt TM (2009) Systematic artifacts in metagenomes from complex microbial communities. The ISME Journal 3(11), 1314–1317.

31. Goñi-Urriza M, Corsellis Y, Lanceleur L, Tessier E, Gury J, Monperrus M, Guyoneaud R (2015) Relationships between bacterial energetic metabolism, mercury methylation potential, and hgcA/hgcB gene expression in *Desulfovibrio dechloroacetivorans* BerOc1. Environmental Science and Pollution Research 22(18), 13764–13771.

32. Handelsman J (2004) Metagenomics: application of genomics to uncultured microorganisms. Microbiology and molecular biology reviews 68(4), 669–685.

33. Haq I, Kumar S, Kumari V, Singh SK, Raj A (2016) Evaluation of bioremediation potentiality of ligninolytic *Serratia liquefaciens* for detoxification of pulp and paper mill effluent. Journal of hazardous materials 305, 190–199.

34. Hess M, Sczyrba A, Egan R, Kim TW, Chokhawala H, Schroth G, Luo S, Clark DS, Chen F, Zhang T, Mackie RI (2011) Metagenomic discovery of biomass-degrading genes and genomes from cow rumen. Science 331(6016), 463–467.

35. Hooper DU, Chapin FS, Ewel JJ, Hector A, Inchausti P, Lavorel S, Lawton JH, Lodge DM, Loreau M, Naeem S, Schmid B (2005) Effects of biodiversity on ecosystem functioning: a consensus of current knowledge. Ecological monographs 75(1), 3–5.

36. Isaac P, Martínez FL, Bourguignon N, Sánchez LA, Ferrero MA (2015) Improved PAHs removal performance by a defined bacterial consortium of indigenous *Pseudomonas* and actinobacteria from Patagonia, Argentina. International Biodeterioration & Biodegradation 101, 23–31.

37. Iverson V, Morris RM, Frazar CD, Berthiaume CT, Morales RL, Armbrust EV (2012) Untangling genomes from metagenomes: revealing an uncultured class of marine Euryarchaeota. Science 335(6068), 587–590.

38. Joutey NT, Bahafid W, Sayel H, Ananou S, El Ghachtouli N (2014) Hexavalent chromium removal by a novel *Serratia proteamaculans* isolated from the bank of Sebou River (Morocco). Environmental Science and Pollution Research 21(4), 3060–3072.

39. Jozefczak M, Remans T, Vangronsveld J, Cuypers A (2012). Glutathione is a key player in metal-induced oxidative stress defenses. International journal of molecular sciences 13(3), 3145–75.

40. Kanehisa M, Sato Y, Morishima K (2016) BlastKOALA and GhostKOALA: KEGG tools for functional characterization of genome and metagenome sequences. Journal of molecular biology 428(4), 726–731.

41. Karnachuk OV, Mardanov AV, Avakyan MR, Kadnikov VV, Vlasova M, Beletsky AV, Gerasimchuk AL, Ravin NV (2015) Draft genome sequence of the first acid-tolerant sulfate-reducing deltaproteobacterium *Desulfovibrio* sp. TomC having potential for minewater treatment. FEMS microbiology letters 362(4), 1–3.

42. Korlevi M, Zucko J, Dragi MN, Blažina M, Pustijanac E, Zeljko TV, Gacesa R, Baranasic D, Starcevic A, Diminic J, Long PF (2015) Bacterial diversity of polluted surface sediments in the northern Adriatic Sea. Systematic and applied microbiology 38(3), 189–197.

43. Labbé D, Margesin R, Schinner F, Whyte LG, Greer CW (2007) Comparative phylogenetic analysis of microbial communities in pristine and hydrocarbon-contaminated Alpine soils. FEMS microbiology ecology 59(2), 466–475.

44. Lavery TJ, Roudnew B, Seymour J, Mitchell JG, Jeffries T (2012) High nutrient transport and cycling potential revealed in the microbial metagenome of Australian sea lion (*Neophoca cinerea*) faeces. PLoS One 7(5), e36478.

45. Leininger S, Urich T, Schloter M, Schwark L, Qi J, Nicol GW, Prosser JI, Schuster SC, Schleper C (2006) Archaea predominate among ammonia-oxidizing prokaryotes in soils. Nature 442(7104), 806–809.

46. Lesser MP (2006) Oxidative stress in marine environments: biochemistry and physiological ecology. Annual Review of Physiology 68, 253–278.

47. Lu J, Jin Q, He Y, He X, Zhao J (2014). Simultaneous removal of phenol and ammonium using Serratia sp. LJ-1 capable of heterotrophic nitrification-aerobic denitrification. Water, Air, & Soil Pollution 225(9), 2125.

48. Lumppio HL, Shenvi NV, Summers AO, Voordouw G, Kurtz DM (2001) Rubrerythrin and Rubredoxin Oxidoreductase in *Desulfovibrio vulgaris*: a Novel Oxidative Stress Protection System. Journal of Bacteriology 183(1), 101–108.

49. Lyons KG, Brigham CA, Traut BH, Schwartz MW (2005) Rare species and ecosystem functioning. Conservation Biology 9(4), 1019–24.

50. MacDougall AS, McCann KS, Gellner G, Turkington R (2013) Diversity loss with persistent human disturbance increases vulnerability to ecosystem collapse. Nature 494(7435), 86–89.

51. Martinez JL (2009) Environmental Pollution by antibiotics and by antibiotic resistance determinants. Environmental pollution 157(11), 2893–902.

52. May RM (1975) Patterns of species abundance and diversity. In: M. L. Cody and J. M. Diamond (ed) Ecology and Evolution of communities, Belnap/Harvard Univ. Press, p 81–120.

53. Meyer F, Paarmann D, D’Souza M, Olson R, Glass EM, Kubal M, Paczian T, Rodriguez A, Stevens R, Wilke A, Wilkening J (2008) The metagenomics RAST server–a public resource for the automatic phylogenetic and functional analysis of metagenomes. BMC bioinformatics 9(1), 386.

54. Mouillot D, Bellwood DR, Baraloto C, Chave J, Galzin R, Harmelin-Vivien M, Kulbicki, M, Lavergne S, Lavorel S, Mouquet N, Paine CT (2013) Rare species support vulnerable functions in high-diversity ecosystems. PLoS biology 11(5), e1001569.

55. Nagayama H, Sugawara T, Endo, R, Ono, A, Kato, H, Ohtsubo Y, Nagata Y, Tsuda M, (2015) Isolation of oxygenase genes for indigo-forming activity from an artificially polluted soil metagenome by functional screening using Pseudomonas putida strains as hosts. Applied microbiology and biotechnology 99(10), 4453–4470.

56. Ntougias S, Lapidus A, Han J, Mavromatis K, Pati A, Chen A, Klenk HP, Woyke T, Fasseas C, Kyrpides NC, Zervakis GI (2014) High quality draft genome sequence of *Olivibacter sitiensis* type strain (AW-6 T), a diphenol degrader with genes involved in the catechol pathway. Standards in genomic sciences 9(3), 783.

57. Olivera ER, Carnicero D, Jodra R, Miñambres B, García B, Abraham GA, Gallardo A, Román JS, García JL, Naharro G, Luengo JM (2001). Genetically engineered *Pseudomonas*: a factory of new bioplastics with broad applications. Environmental microbiology 3(10), 612–618.

58. Overbeek R, Begley T, Butler RM, Choudhuri JV, Chuang HY, Cohoon M, de Crécy-Lagard V, Diaz N, Disz T, Edwards R, Fonstein M (2005) The subsystems approach to genome annotation and its use in the project to annotate 1000 genomes. acids research 33(17), 5691- 5702.

59. Parks DH, Tyson GW, Hugenholtz P, Beiko RG (2014) STAMP: statistical analysis of taxonomic and functional profiles. Bioinformatics 30(21), 3123–3124.

60. Patel V, Munot H, Shouche YS, Madamwar D (2014) Response of bacterial community structure to seasonal fluctuation and anthropogenic pollution on coastal water of Alang– Sosiya ship breaking yard, Bhavnagar, India. Bioresource technology 161, 362–370.

61. Pat-Espadas AM, Razo-Flores, E, Rangel-Mendez JR, Cervantes FJ (2013) Reduction of palladium and production of nano-catalyst by *Geobacter sulfurreducens*. Applied microbiology and biotechnology 97(21), 9553–9560.

62. Payne RB, Gentry DM, Rapp-Giles BJ, Casalot L, Wall JD (2002) Uranium reduction by *Desulfovibrio desulfuricans* strain G20 and a cytochrome c3 mutant. Applied and Environmental Microbiology 68(6), 3129–3132.

63. Pieper DH, Reineke W (2000) Engineering bacteria for bioremediation. Current opinion in biotechnology 11(3), 262–270.

64. Prakash OM, Gihring TM, Dalton, D.D., Chin, K.J., Green, S.J., Akob, D.M., Wanger G., Kostka JE (2010) *Geobacter daltonii* sp. nov., an Fe (III)-and uranium (VI)-reducing bacterium isolated from a shallow subsurface exposed to mixed heavy metal and hydrocarbon contamination. International journal of systematic and evolutionary microbiology 60(3), 546–553.

65. Preston EW (1962) The canonical distribution of commonness and rarity. Part I Ecology. 43, 185–215.

66. Rho M, Tang H, Ye Y (2010) FragGeneScan: predicting genes in short and error-prone reads. Nucleic acids research 38(20), e191.

67. Romic M, Romic D (2003) Heavy metals distribution in agricultural topsoils in urban area. Environmental geology 43(7), 795–805.

68. Sangwan N, Lata P, Dwivedi V, Singh A, Niharika N, Kaur J, Anand S, Malhotra J, Jindal S, Nigam A, Lal R (2012) Comparative metagenomic analysis of soil microbial communities across three hexachlorocyclohexane contamination levels. PLoS One 7(9), e46219.

69. Schmieder R, Edwards R (2011) Quality control and preprocessing of metagenomic datasets. Bioinformatics 27(6), 863–864.

70. Shah V, Zakrzewski M, Wibberg D, Eikmeyer F, Schlüter A, Madamwar D (2013) Taxonomic profiling and metagenome analysis of a microbial community from a habitat contaminated with industrial discharges. Microbial ecology 66(3), 533–550.

71. Sharaff M, Kamat S, Archana G (2017) Analysis of copper tolerant rhizobacteria from the industrial belt of Gujarat, western India for plant growth promotion in metal polluted agriculture soils. Ecotoxicology and environmental safety 138, 113–121.

72. Shokralla S, Spall JL, Gibson JF, Hajibabaei M (2012) Next-generation sequencing technologies for environmental DNA research. Molecular ecology 21(8), 1794–1805.

73. Singh A, Chauhan NS, Thulasiram HV, Taneja V, Sharma R (2010) Identification of two flavin monooxygenases from an effluent treatment plant sludge metagenomic library. Bioresource technology 101(21), 8481–8484.

74. Singleton DR, Ramirez LG, Aitken MD (2009) Characterization of a polycyclic aromatic hydrocarbon degradation gene cluster in a phenanthrene-degrading *Acidovorax* strain. Applied and environmental microbiology 75(9), 2613–2620.

75. Smith MD, Knapp AK (2003). Dominant species maintain ecosystem function with non-random species loss. Ecology Letters 6(6), 509–517.

76. Sukul P, Spiteller M (2007) Fluoroquinolone antibiotics in the environment. In Reviews of environmental contamination and toxicology pp. 131–162.

77. Sutton NB, Maphosa F, Morillo JA, Al-Soud WA, Langenhoff AA, Grotenhuis T, Rijnaarts HH, Smidt H (2013) Impact of long-term diesel contamination on soil microbial community structure. Applied and environmental microbiology 79(2), 619–630.

78. Tokeshi M (1993) Species abundance patterns and community structure. Advances in ecological research 24, 111–186.

79. Torrentó C, Cama J, Urmeneta J, Otero N, Soler A (2010) Denitrification of groundwater with pyrite and *Thiobacillus denitrificans*. Chemical Geology 278(1), 80–91.

80. Ufarté L, Laville É, Duquesne S, Potocki-Veronese G (2015) Metagenomics for the discovery of pollutant degrading enzymes. Biotechnology advances 33(8), 1845–1854.

81. Varin T, Lovejoy C, Jungblut AD, Vincent WF, Corbeil J (2012) Metagenomic analysis of stress genes in microbial mat communities from Antarctica and the High Arctic. Applied and environmental microbiology 78(2), 549–559.

82. Webber MA, Piddock LJ (2003). The importance of efflux pumps in bacterial antibiotic resistance. Journal of Antimicrobial Chemotherapy 51(1), 9–11.

83. Wickramasekara S, Neilson J, Patel N, Breci L, Hilderbrand A, Maier RM, Wysocki V (2011) Proteomics analyses of the opportunistic pathogen *Burkholderia vietnamiensis* using protein fractionations and mass spectrometry. BioMed Research International. Article ID 701928, doi:10.1155/2011/701928

84. Xu M, He Z, Zhang Q, Liu J, Guo J, Sun G, Zhou J (2015) Responses of aromatic-degrading microbial communities to elevated nitrate in sediments. Environmental science & technology 49(20), 12422–12431.

85. Yang S, Wen X, Jin, H, Wu Q (2012) Pyrosequencing investigation into the bacterial community in permafrost soils along the China-Russia Crude Oil Pipeline (CRCOP). PLoS One 7(12), e52730.

86. Zhang T, Shao MF, Ye L (2012) 454 Pyrosequencing reveals bacterial diversity of activated sludge from 14 sewage treatment plants. The ISME journal 6(6), 1137–1147.

87. Zhou S, Xu J, Yang G, Zhuang L (2014) Methanogenesis affected by the co-occurrence of iron (III) oxides and humic substances. FEMS microbiology ecology 88(1), 107–120.

